# Predicting resistance of clinical Abl mutations to targeted kinase inhibitors using alchemical free-energy calculations

**DOI:** 10.1101/239012

**Authors:** Kevin Hauser, Christopher Negron, Steven K. Albanese, Soumya Ray, Thomas Steinbrecher, Robert Abel, John D. Chodera, Lingle Wang

## Abstract

The therapeutic effect of targeted kinase inhibitors can be significantly reduced by intrinsic or acquired resistance mutations that modulate the affinity of the drug for the kinase. In cancer, the majority of missense mutations are rare, making it difficult to predict their impact on inhibitor affinity. This complicates the practice of precision medicine, pairing of patients with clinical trials, and development of next-generation inhibitors. Here, we examine the potential for alchemical free-energy calculations to predict how kinase mutations modulate inhibitor affinities to Abl, a major target in chronic myelogenous leukemia (CML). We find these calculations can achieve useful accuracy in predicting resistance for a set of eight FDA-approved kinase inhibitors across 144 clinically-identified point mutations, achieving a root mean square error in binding free energy changes of 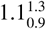 kcal/mol (95% confidence interval) and correctly classifying mutations as resistant or susceptible with 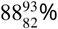 accuracy. Since these calculations are fast on modern GPUs, this benchmark establishes the potential for physical modeling to collaboratively support the rapid assessment and anticipation of the potential for patient mutations to affect drug potency in clinical applications.

---

Targeted kinase inhibitors are a major therapeutic class in the treatment of cancer. A total of 38 selective small molecule kinase inhibitors have now been approved by the FDA [1], including 34 approved to treat cancer, and perhaps 50% of all current drugs in development target kinases [2]. Despite the success of selective inhibitors, the emergence of drug resistance remains a challenge in the treatment of cancer [3–10] and has motivated the development of second-and then third-generation inhibitors aimed at overcoming recurrent resistance mutations [11–15].

While a number of drug resistance mechanisms have been identified in cancer (e.g., induction of splice variants [16], or alleviation of feedback [17]), inherent or acquired missense mutations in the kinase domain of the target of therapy are a major form of resistance to tyrosine kinase inhibitors (TKI) [10, 18, 19]. Oncology is entering a new era with major cancer centers now deep sequencing tumors to reveal genetic alterations that may render subclonal populations susceptible or resistant to targeted inhibitors [20], but the use of this information in precision medicine has lagged behind. It would be of enormous value in clinical practice if an oncologist could reliably ascertain whether these mutations render the target of therapy resistant or susceptible to available inhibitors; such tools would facilitate the enrollment of patients in mechanism-based basket trials [21, 22], help prioritize candidate compounds for clinical trials, and aid the development of next-generation inhibitors.

## The long tail of rare kinase mutations frustrates prediction of drug resistance

While some cancer missense mutations are highly recurrent and have been characterized clinically or biochemically, a “long tail” of rare mutations collectively accounts for the majority of clinically observed missense mutations (***Figure 1***a), leaving clinicians and researchers without knowledge of whether these uncharacterized mutations might lead to resistance. While rules-based and machine learning schemes are still being assessed in oncology contexts, work in predicting drug response to microbial resistance has shown that rare mutations present a significant challenge to approaches that seek to predict resistance to therapy [23]. Clinical cancer mutations may impact drug response through a variety of mechanisms by altering kinase activity, ATP affinity, substrate specificities, and the ability to participate in regulatory interactions, compounding the difficulties associated with limited datasets that machine learning approaches face. In parallel with computational approaches, high-throughput experimental techniques such as MITE-Seq [24] have been developed to assess the impact of point mutations on drug response. However, the complexity of defining selection schemes that reliably correlate with *in vivo* drug effectiveness and long turn-around times might limit their ability to rapidly and reliably impact clinical decision-making.

**Figure 1.**
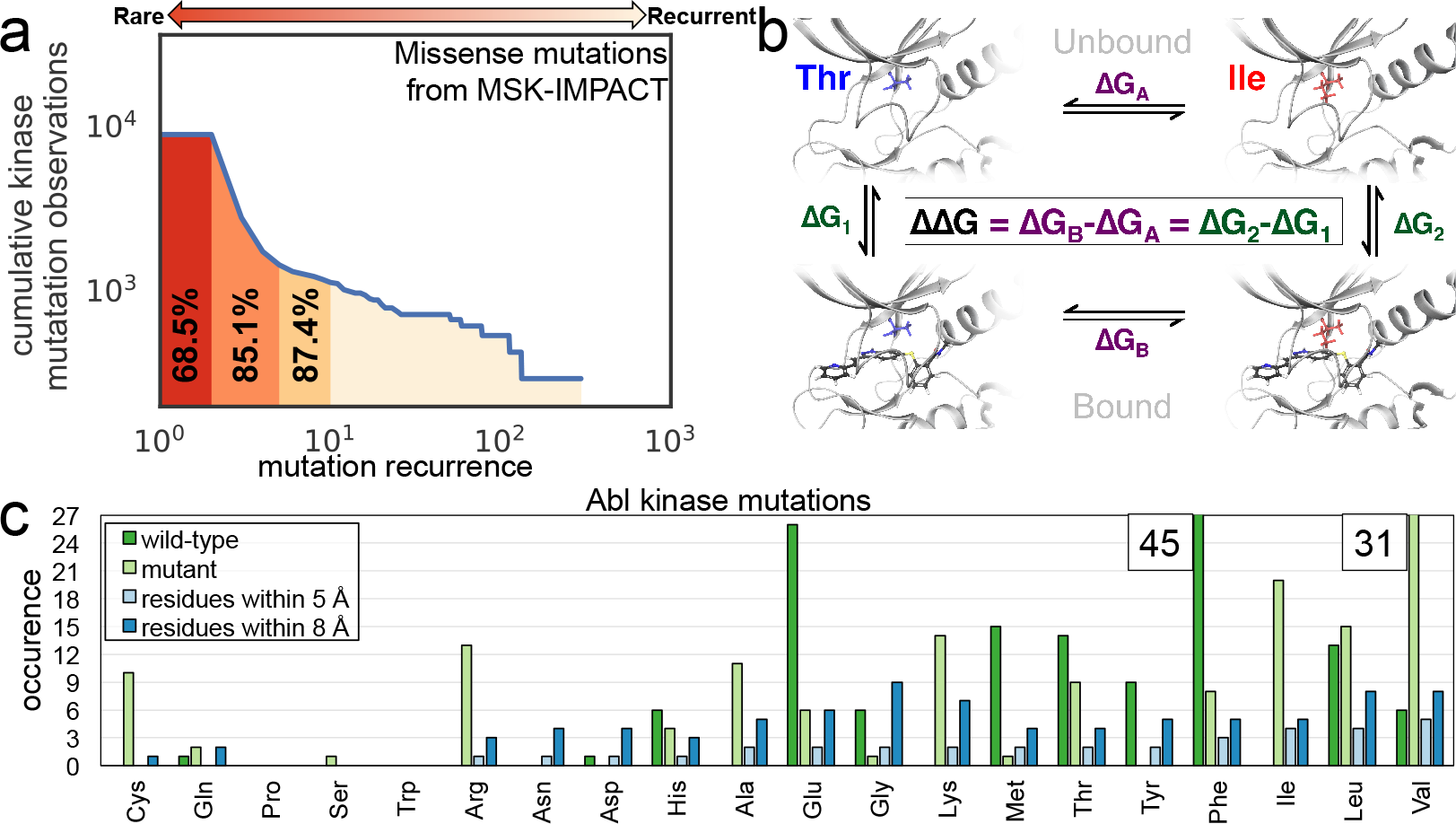
Relative alchemical free-energy calculations can be used to predict affinity changes of FDA-approved selective kinase inhibitors arising from clinically-identified mutations in their targets of therapy. (*a*) Missense mutation statistics derived from 10,336 patient samples subjected to MSK-IMPACT deep sequencing panel [20] show that 68.5% of missense kinase mutations in cancer patients have never been observed previously, while 87.4% have been observed no more than ten times. (*b*) To compute the impact of a clinical point mutation on inhibitor binding free energy, a thermodynamic cycle can be used to relate the free energy of the wild-type and mutant kinase in the absence (top) and presence (bottom) of the inhibitor. (*c*) Summary of mutations studied in this work. Frequency of the wild-type (dark green) and mutant (green) residues for the 144 clinically-identified Abl mutations used in this study (see ***Table 1*** for data sources). Also shown is the frequency of residues within 5 Å (light blue) and 8 Å (blue) of the binding pocket. The number of wild-type Phe residues (n=45) and mutant Val residues (n=31) exceeded the limits of the y-axis.

## Alchemical free-energy methods can predict inhibitor binding affinities

Physics-based approaches could be complementary to machine-learning and experimental techniques in predicting changes in TKI affinity due to mutations with few or no prior clinical observations. Modern atomistic molecular mechanics forcefields such as OPLS3 [25], CHARMM [26], and AMBER FF14SB [27] have reached a sufficient level of maturity to enable the accurate and reliable prediction of receptor-ligand binding free energy. Alchemical free-energy methods permit receptor-ligand binding energies to be computed rigorously, including all relevant entropic and enthalpic contributions [28]. Encouragingly, kinase:inhibitor binding affinities have been predicted using alchemical free-energy methods with mean unsigned errors of 1.0 kcal/mol forCDK2,JNK1, p38, and Tyk2 [29, 30]. Beyond kinases, alchemical approaches have predicted the binding affinity of BRD4 inhibitors with mean absolute errors of 0.6 kcal/mol [31]. Alchemical methods have also been observed to have good accuracy (0.6 kcal/mol mean unsigned error for Tyk2 tyrosine kinase) in the prediction of relative free energies for ligand transformations within a complex whose receptor geometry was generated using a homology model [32].

## Alchemical approaches can predict the impact of protein mutations on free energy

Alchemical free-energy calculations have also been used to predict the impact of mutations on protein-protein binding [33] and protein thermostabilities [34]. Recent work has found that protein mutations can be predicted to be stabilizing or destabilizing with a classification accuracy of 71% across ten proteins and 62 mutations [35]. The impact of Gly to D-Ala mutations on protein stability was predicted using an alchemical approach with a similar level of accuracy [36]. Recently, one study has hinted at the potential utility of alchemical free-energy calculations in oncology by predicting the impact of a single clinical mutation on the binding free energies of the TKIs dasatinib and RL45 [37].

## Assessing the potential for physical modeling to predict resistance to FDA-approved TKIs

Here, we ask whether physical modeling techniques may be useful in predicting whether clinically-identified kinase mutations lead to drug resistance or drug sensitivity. We perform state-of-the-art relative alchemical free-energy calculations using FEP+ [29], recently demonstrated to achieve sufficiently good accuracy to drive the design of small-molecule inhibitors for a broad range of targets during lead optimization [28–30, 38]. We compare this approach against a fast but approximate physical modeling method implemented in Prime [39] (an MM-GBSA approach) in which an implicit solvent model is used to assess the change in minimized interaction energy of the ligand with the mutant and wild-type kinase. We consider whether these methods can predict a ten-fold reduction in inhibitor affinity (corresponding to a binding free energy change of 1.36 kcal/mol) to assess baseline utility. As a benchmark, we compile a set of reliable inhibitor ΔpIC_50_ data for 144 clinically-identified mutants of the human kinase Abl, an important oncology target dysregulated in cancers like chronic myelogenous leukemia (CML), for which six [1] FDA-approved TKIs are available. While ΔpIC_50_ can approximate a dissociation constant Δ*K_D_*, other processes contributing to changes in cell viability might affect IC_50_ in ways that are not accounted for by a traditional binding experiment, motivating a quantitative comparison between ΔpIC_50_ and Δ*K_D_*. The results of this benchmark demonstrate the potential for FEP+ to predict the impact that mutations in Abl kinase have on drug binding, and a classification accuracy of 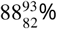 (for all statistical metrics reported in this paper, the 95% confidence intervals (CI) is shown in the form of 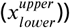, an RMSE of 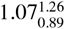 kcal/mol, and an MUE of 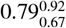 kcal/mol was achieved.

## Results

### Free energy calculations can recapitulate the impact of clinical mutations on TKI affinity

Alchemical free-energy calculations utilize a physics-based approach to estimate the free energy of transforming one chemical species into another, incorporating all enthalpic and entropic contributions in a physically consistent manner [28, 40–42]. While relative alchemical free-energy calculations have typically been employed in optimizing small molecules for increased potency or selectivity [29, 38, 42, 43], a complementary alchemical approach can be used to compute the impact of point mutations on ligand binding affinities. ***Figure 1***b depicts the thermodynamic cycle that illustrates how we used relative free energy calculations to compute the change in ligand binding free energy in response to the introduction of a point mutation in the kinase. In the *bound* leg of the cycle, the wild-type protein:ligand complex is transformed into the mutant protein:ligand complex. In the *unbound* leg of the cycle, the *apo* protein is transformed from wild-type into mutant. To achieve reliable predictions with short relative free-energy calculations, a reliable receptor:ligand complex structure is required with the assumption that the binding mode of wild-type and mutant are similar. In this work, high-resolution co-crystal structures of wild-type Abl bound to an inhibitor were utilized when available. To assess the potential for using docked inhibitor poses, we also examined two systems for which co-crystal structures were not available (Abl:erlotinib and Abl:gefitinib) and used docking to generate initial coordinates.

### Compiled ΔpIC_50_ data provides a benchmark for predicting mutational resistance

To construct a benchmark evaluation dataset, we compiled a total of 144 ΔpIC_50_ measurements of Abl:TKI affinities, summarized in ***Table 1***, taking care to ensure all measurements for an individual TKI were reported in the same study from experiments run under identical conditions. 131 ΔpIC_50_ measurements were available across the six TKIs with available co-crystal structures with wild-type Abl—26 for axitinib and 21 for bosutinib, dasatinib, imatinib, nilotinib, and ponatinib. 13 ΔpIC_50_ measurements were available for the two TKIs for which docking was necessary to generate Abl:TKI structures—7 for erlotinib and 6 for gefitinib. For added diversity, this set includes TKIs for which Abl is not the primary target—axitinib, erlotinib, and gefitinib. All mutations in this benchmark dataset have been clinically-observed (***Table S1***). Due to the change in bond topology required by mutations involving proline, which is not currently supported by the FEP+ technology for protein residue mutations, the three mutations H396P (axitinib, gefitinib, erlotinib) were excluded from our assessment. As single point mutations were highly represented in the IMPACT study analyzed in ***Figure 1***a, we excluded double mutations from this work. However, the impact of mutations from multiple sites can potentially be modeled by sequentially mutating each site and this will be addressed in future work.

**Table 1.**
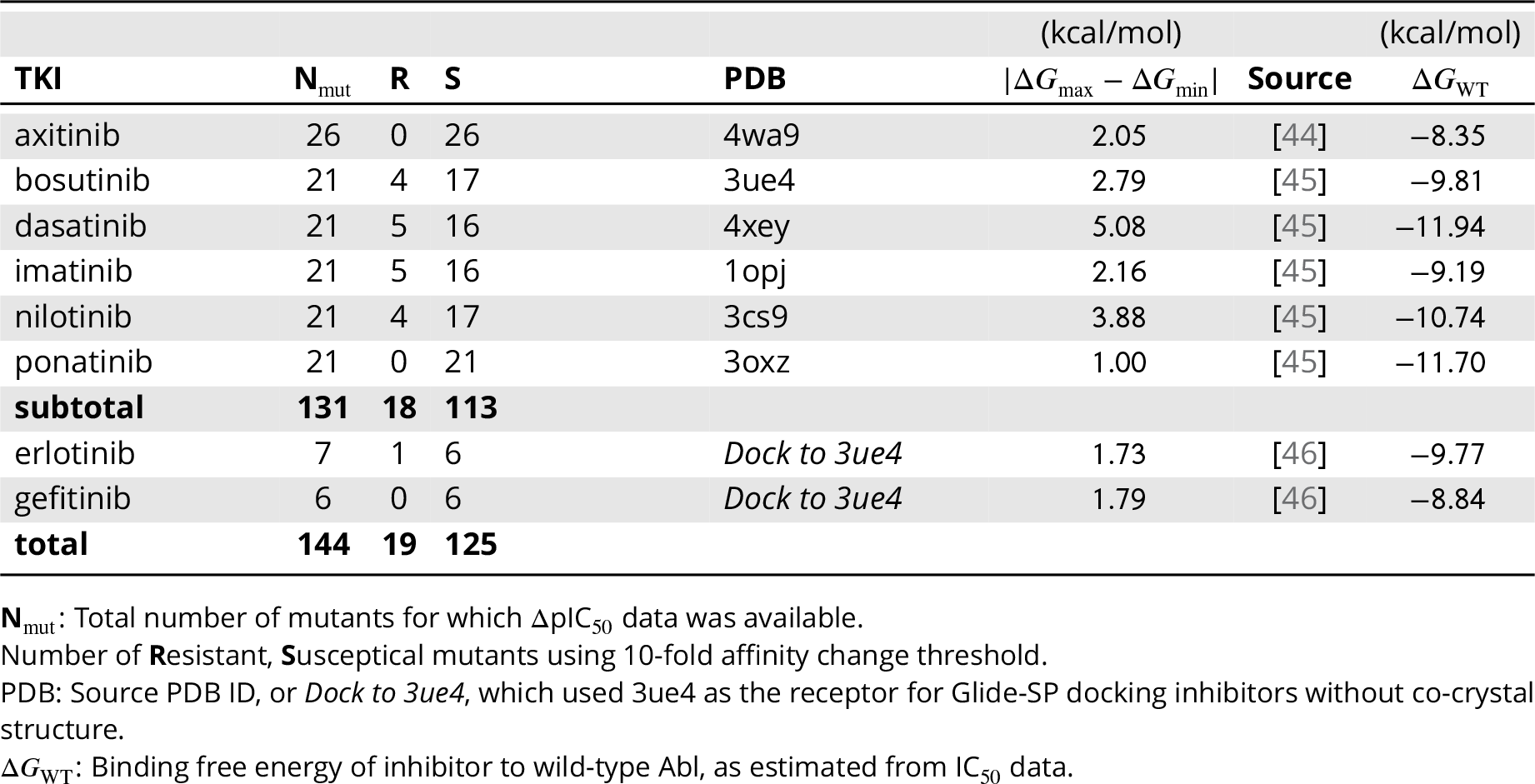
Public ΔpIC_50_ datasets for 144 Abl kinase mutations and eight tyrosine kinase inhibitors (TKIs) with corresponding wild-type co-crystal structures used in this study.

| TKI | $N_{mut}$ | R | S | PDB | (kcal/mol) | Source | (kcal/mol) |
| --- | --- | --- | --- | --- | --- | --- | --- |
| | | | | | $\vert \Delta G_{max} - \Delta G_{min}\vert $ | | $\Delta G_{WT}$ |
| axitinib | 26 | 0 | 26 | 4wa9 | 2.05 | [44] | -8.35 |
| bosutinib | 21 | 4 | 17 | 3ue4 | 2.79 | [45] | -9.81 |
| dasatinib | 21 | 5 | 16 | 4xey | 5.08 | [45] | -11.94 |
| imatinib | 21 | 5 | 16 | 1opj | 2.16 | [45] | -9.19 |
| nilotinib | 21 | 4 | 17 | 3cs9 | 3.88 | [45] | -10.74 |
| ponatinib | 21 | 0 | 21 | 3oxz | 1.00 | [45] | -11.70 |
| **subtotal** | **131** | **18** | **113** |  |  |  |  |
| erlotinib | 7 | 1 | 6 | *Dock to 3ue4* | 1.73 | [46] | -9.77 |
| gefitinib | 6 | 0 | 6 | *Dock to 3ue4* | 1.79 | [46] | -8.84 |
| **total** | **144** | **19** | **125** |  |  |  |  |

$N_{mut}$ : Total number of mutants for which $\Delta pIC_{50}$ data was available.
Number of **R**esistant, **S**usceptible mutants using 10-fold affinity change threshold.
PDB: Source PDB ID, or *Dock to 3ue4*, which used 3ue4 as the receptor for Glide-SP docking inhibitors without co-crystal structure.
$\Delta G_{WT}$ : Binding free energy of inhibitor to wild-type Abl, as estimated from $IC_{50}$ data.

Experimental ΔpIC_50_ measurements for wild-type and mutant Abl were converted to ΔΔG in order to make direct comparisons between physics-based models and experiment. However, computation of experimental uncertainties were required to understand the degree to which differences between predictions and experimental data were significant. Since experimental error estimates for measured IC_50_s were not available for the data in ***Table 1***, we compared that data to other sources that have published IC_50_s for the same mutations in the presence of the same TKIs (***Figure 2***a,b,c). Cross-comparison of 97 experimentally measured ΔΔGs derived from cell viability assay IC_50_ data led to an estimate of experimental variability of 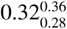 kcal/mol root-mean square error (RMSE) that described the expected repeatability of the measurements. Because multiple factors influence the IC_50_ aside from direct effects on the binding affinity—the focus of this study—we also compared ΔΔGs derived from ΔpIC_50_s with those derived from binding affinity measurements (Δ*K*_*d*_) for which data for a limited set of 27 mutations was available (***Figure 2***d); the larger computed RMSE of 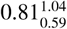 kcal/mol represents an estimate of the lower bound of the RMSE to the IC_50_-derived ΔΔGs that we might hope to achieve with FEP+ or Prime, which were performed using non-phosphorylated models, when comparing sample statistics directly. In comparing 31 mutations for which phosphorylated and non-phosphorylated Δ*K_d_*s were available, we found a strong correlation between the ΔΔGs derived from those data (r=0.94, Supplementary ***Figure S1***); the statistics of that comparison are similar to those of the inter-lab variability comparison.

**Figure 2.**
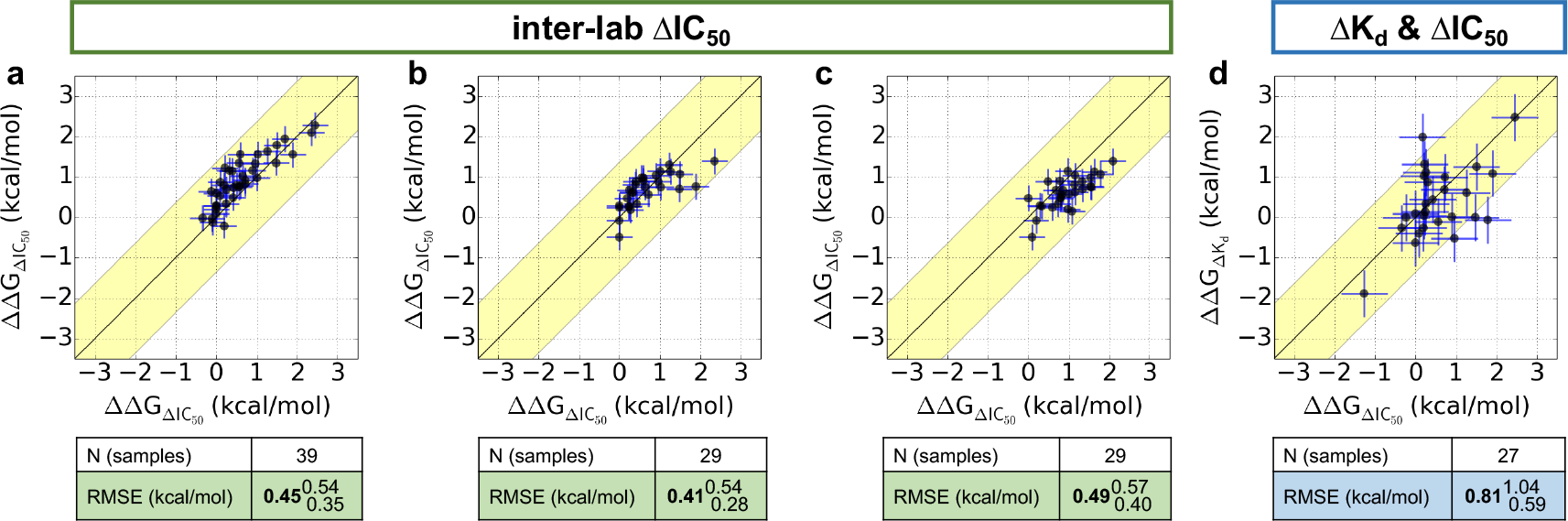
Cross-comparison of the experimentally measured effects that mutations in Abl kinase have on ligand binding, performed by different labs. ΔΔG was computed from publicly available ΔpIC_50_ or Δp*K_d_* measurements and these values of ΔΔG were then plotted and the RMSE between them reported. (**a**) ΔpIC_50_ measurements (X-axis) from [45] compared with ΔpIC_50_ measurements (Y-axis) from [47]. (**b**) ΔpIC_50_ measurements (X-axis) from [45] compared with ΔpIC_50_ measurements (Y-axis) from [48]. (**c**) ΔpIC_50_ measurements (X-axis) from [47] compared with ΔpIC_50_ measurements (Y-axis) from [48]. (**d**) ΔpIC_50_ measurements (X-axis) from [45] compared with Δp*K_d_* measurements (Y-axis) from [46] using non-phosphorylated Abl kinase. Scatter plot error bars in a, b, and c are ±standard error (SE) taken from the combined 97 inter-lab ΔΔGs derived from the ΔpIC_50_ measurements, which was 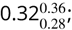 the RMSE was 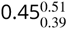 kcal/mol. Scatter plot error bars in d are the ±standard error (SE) of ΔΔGs derived from ΔpIC_50_ and Δp*K_d_* from a set of 27 mutations, which is 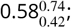 the RMSE was 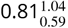 kcal/mol.

### Most clinical mutations do not significantly reduce TKI potency

The majority of mutations do not lead to resistance by our 10-fold affinity loss threshold: 86.3% of the co-crystal set(n=113) and 86.8% of the total set(n=125). Resistance mutations, which are likely to result in a failure of therapy, constitute 13.7% of the co-crystal set(n=18) and 13.2% of the total set of mutations (n=19). The ΔpIC_50_s for all 144 mutations are summarized in ***Table S2***—***Table S7*** in the Supplementary Information. Two mutations exceeded the dynamic range of the assays (IC_50_ >10,000 nM); as these two mutations clearly raise resistance, we excluded them from quantitative analysis (RMSE and MUE) but included them in truth table analyses and classification metrics (accuracy, specificity, sensitivity).

### How accurately does physical modeling predict affinity changes for clinical Abl mutants?

From prior experience with relative alchemical free-energy calculations for ligand design, good initial receptor-ligand geometry was critical to obtaining accurate and reliable free energy predictions [29], so we first focused on the 131 mutations in Abl kinase across six TKIs for which wild-type Abl:TKI co-crystal structures were available. ***Figure 3*** summarizes the performance of predicted binding free-energy changes (ΔΔG) for all 131 mutants in this set for both a fast MM-GBSA physics-based method that only captures interaction energies for a single structure (Prime) and rigorous alchemical free-energy calculations (FEP+). Scatter plots compare experimental and predicted free-energy changes (ΔΔG) and characterize the ability of these two techniques to predict experimental measurements. Statistical uncertainty in the predictions and experiment-to-experiment variability in the experimental values are shown as ellipse height and widths respectively. The value for experimental variability was 0.32 kcal/mol, which was the standard error computed from the cross-comparison in ***Figure 2***. For FEP+, the uncertainty was taken to be the standard error of the average from three independent runs for a particular mutation, while Prime results are deterministic and are not contaminated by statistical uncertainty (see Methods).

To better assess whether discrepancies between experimental and computed ΔΔ*G*s simply arise for known forcefield limitations or might indicate more significant effects, we incorporated an additional error model in which the forcefield error was taken to be a random error *σ*_FF_ ≈ 0.9 kcal/mol, a value established form previous benchmarks on small molecules absent conformational sampling or protonation state issues [25]. Thin error bars in ***Figure 2*** represent the overall estimated error due to both this forcefield error and experimental variability or statistical uncertainty.

To assess overall quantitative accuracy, we computed both root-mean-squared error (RMSE)—which is rather sensitive to outliers, and mean unsigned error (MUE). For Prime, the MUE was 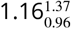 kcal/mol and the RMSE was 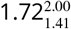 kcal/mol. FEP+, the alchemical free-energy approach, achieved a significantly higher level of quantitative accuracy with an MUE of 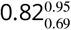 kcal/mol and an RMSE of 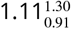 kcal/mol. Notably, alchemical free energy calculations come substantially closer than MMGBSA approach to the minimum achievable RMSE of 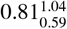 kcal/mol (due to experimental error; ***Figure 2***) for this dataset.

### How accurately can physical modeling classify mutations as susceptible or resistant?

While quantitative accuracy (MUE, RMSE) is a principle metric of model performance, an application of potential interest is the ability to classify mutations as causing resistance to a specific TKI. To characterize the accuracy with which Prime and FEP+ classified mutations in a manner that might be therapeutically relevant, we classified mutations by their experimental impact on the binding affinity as *susceptible* (affinity for mutant is diminished by no more than 10-fold, ΔΔ*G* ≤ 1.36 kcal/mol) or as *resistant* (affinity for mutant is diminished by least 10-fold, ΔΔ*G* > 1.36 kcal/mol). Summary statistics of experimental and computational predictions of these classes are shown in ***Figure 2*** (bottom) as truth tables (also known as *confusion matrices*).

The simple minimum-energy scoring method Prime correctly classified 9 of the 18 resistance mutations in the dataset while merely 85 of the 113 susceptible mutations were correctly classified (28 false positives). In comparison, the alchemical free-energy method FEP+, which includes entropic and enthalpic contributions as well as explicit representation of solvent, correctly classified 9 of the 18 resistance mutations while a vast majority, 105, of the susceptible mutations were correctly classified (merely 8 false positives). Prime achieved a classification accuracy of 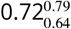, while FEP+ achieved an accuracy that is significantly higher (both in a statistical sense and in overall magnitude), achieving an accuracy of 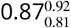. Sensitivity (also called *true positive rate*) and specificity (*true negative rate*) are also informative statistics in assessing the performance of a binary classification scheme. For Prime, the sensitivity was 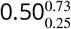, while the specificity was 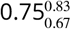. To put this in perspective, a CML patient bearing a resistance mutation in the kinase domain of Abl has an equal chance of Prime correctly predicting this mutation would be resistant to one of the TKIs considered here, while if the mutation was susceptible, the chance of correct prediction would be ~75%. By contrast, the classification specificity of FEP+ was substantially better. For FEP+, the sensitivity was 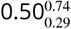 while the specificity was 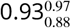. There is a very high probability that FEP+ will correctly predict that one of the eight TKIs studied here will remain effective for a patient bearing a susceptible mutation.

### How sensitive are classification results to choice of cutoff?

Previous work by O’Hare et al. utilized TKI-specific thresholds for dasatinib, imatinib, and nilotinib [49], which were ~2 kcal/mol. Supplementary ***Figure S2*** shows that when our classification threshold was increased to a 20-fold change in binding (1.77 kcal/mol), FEP+ correctly classified 8 of the 13 resistant mutations and with a threshold of 100-fold change in binding (2.72 kcal/mol), FEP+ correctly classified the only two resistant mutations (T315I/dasatinib and T315I/nilotinib). With the extant multilayered and multinodal decision-making algorithms used by experienced oncologists to manage their patients’ treatment, or by medicinal chemists to propose candidate compounds for clinical trials, the resistant or susceptible cutoffs could be selected with more nuance than the simple 10-fold affinity threshold we consider here. With a larger affinity change cutoff, for example, the accuracy with which physical models predict resistance mutations increases beyond 90% (Supplementary ***Figure S2***). For the alchemical approach, the two-class accuracy was 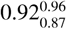 when an affinity change cutoff of 20-fold was used while using an affinity change cutoff of 100-fold further improved the two-class accuracy to 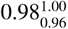.

### Bayesian analysis can estimate the true error

The statistical metrics—MUE, RMSE, accuracy, specificity, and sensitivity—discussed above are based on analysis of the apparent performance of the observed modeling results compared with the observed experimental data via sample statistics. However, this analysis considers a limited number of mutants, and both measurements and computed values are contaminated with experimental or statistical error. To obtain an estimate of the *intrinsic performance* of our physical modeling approaches, accounting for known properties of the experimental variability and statistical uncertainties, we used a hierarchical Bayesian model (detailed in the Methods) to infer posterior predictive distributions from which expectations and 95% predictive intervals could be obtained. The results of this analysis are presented in ***Figure 3*** (central tables).

FEP+ is significantly better than Prime at predicting the impact of mutations on TKI binding affinities, as the apparent performance (using the original observations) as well as the intrinsic performance (where Bayesian analysis was used to correct for statistical uncertainty or experimental variation) were well-separated outside their 95% confidence intervals in nearly all metrics. Applying the Bayesian model, the MUE and RMSE for FEP+ was 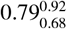 kcal/mol and 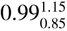 kcal/mol respectively (N=129). For the classification metrics accuracy, specificity, and sensitivity, the model yields 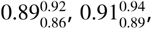 and 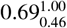 respectively (N=131). The intrinsic RMSE and MUE of Prime was 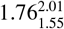 kcal/mol and 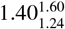 kcal/mol (N=129) respectively, and the classification accuracy, specificity, and sensitivity was 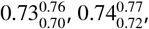 and 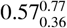 respectively (N=131). The intrinsic MUE of Prime obtained by this analysis is larger than the observed MUE reflecting the non-Gaussian, fat-tailed error distributions of Prime results.

**Figure 3.**
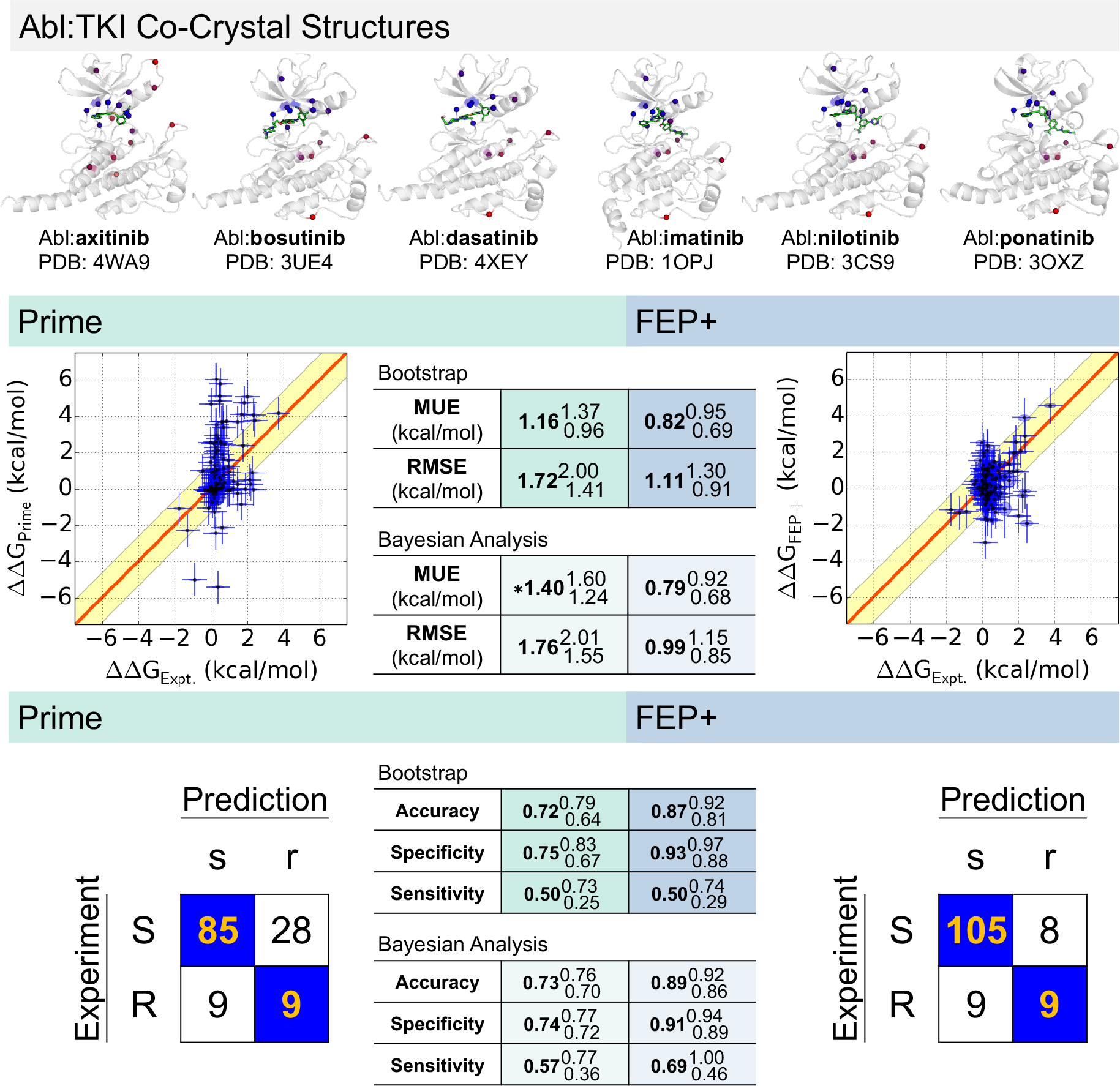
Comparison of experimentally-measured binding free-energy changes (ΔΔG) for 131 clinically observed mutations and 6 selective kinase inhibitors for which co-crystal structures of wild-type kinase with inhibitor are available. *Top panel:* Abl:TKI co-crystal structures used in this study with locations of clinical mutants for each inhibitor highlighted (colored from blue to red for residues nearest to farthest from ligand) in relation to TKI (green sticks) on the corresponding Abl:TKI wild-type crystal structure. *Middle panel:* Scatter plots show Prime and FEP+ computed ΔΔG compared to experiment, with ellipse widths and heights (*±*σ) for experiment and FEP+ respectively. The red diagonal line indicates when prediction equals experiment, while the yellow shaded region indicates area in which predicted ΔΔG is within 1.36 kcal/mol of experiment (corresponding to a ten-fold error in predicted affinity change). ΔΔG < 0 denotes the mutation increases the susceptibility of the kinase to the inhibitor, while ΔΔG > 0 denotes the mutation increases the resistance of the kinase to the inhibitor. The two mutations that were beyond the concentration limit of the assay (T315I/dasatinib, L248R/imatinib) were not plotted; 129 points were plotted. Truth tables of classification accuracy, sensitivity and specificity using two-classes. *Bottom panel:* Truth tables and classification results include T315l/dasatinib and L248R/imatinib; 131 points were used. For MUE, RMSE, and truth table performance statistics, sub/superscripts denote 95 % CIs. Variability in the experimental data is shown as ellipse widths and uncertainty in our calculations is shown as ellipse heights. Experimental variability was computed as the standard error between IC_50_-derived ΔΔG measurements made by different labs, 0.32 kcal/mol. The statistical uncertainty in the Prime calculations was zero because the method is deterministic (σ_cal_ = 0), while the uncertainty in the FEP+ calculations was reported as the standard error, σ_cal_, of the mean of the predicted ΔΔ*G*s from three independent runs. To better highlight true outliers unlikely to simply result from expected forcefield error, we presume forcefield error (σ_FF_ ≈ 0.9 kcal/mol [25]) also behaves as a random error, and represent the total estimated statistical and forcefield error 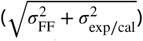 as vertical error bars. The horizontal error bars for the experiment (σ_exp_) was computed as the standard error between ΔpIC_50_ and ΔK_*d*_ measurements, 0.58 kcal/mol. For Prime, **\*MUE** highlights that the Bayesian model yields a value for MUE that is noticeably larger than MUE for observed data due to the non-Gaussian error distribution of Prime.

### Is the impact of point mutations on drug binding equally well-predicted for the six TKIs?

The impact of point mutations on drug binding are not equally well predicted for the six TKIs. ***Figure 4*** expands the results in ***Figure 3*** on a TKI-by-TKI basis to dissect the particular mutations in the presence of a specific TKI. Prime and FEP+ correctly predicted that most mutations in this dataset (N=26) do not raise resistance to axitinib, though FEP+ predicted 4 false positives compared with 3 false positives by Prime. The MUE and RMSE of FEP+ was excellent for this inhibitor, 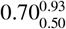 kcal/mol and 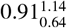 kcal/mol respectively. While the classification results for bosutinib (N=21) were equally well predicted by Prime as by FEP+, FEP+ was still able to achieve superior, but not highly significant, predictive performance for the quantitative metrics MUE and RMSE, which were 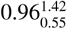 kcal/mol and 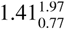 kcal/mol respectively (FEP+) and 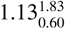 kcal/mol and 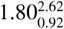 kcal/mol respectively (Prime). For dasatinib, FEP+ achieved an MUE and RMSE of 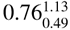 kcal/mol and 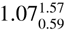 kcal/mol respectively whereas the results were, as expected, less quantitatively predictive for Prime (N=20). The results for imatinib were similar to those of dasatinib above, where the MUE and RMSE for FEP+ were 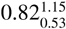 kcal/mol and 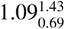 kcal/mol respectively (N=20). Nilotinib, a derivative of imatinib, led to nearly identical quantitative performance results for FEP+ with an MUE and RMSE of 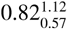 kcal/mol and 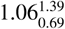 kcal/mol respectively (N=21). Similar to axitinib, ponatinib presented an interesting case because there were no mutations in this dataset that raised resistance. Despite the wide dynamic range in the computed values of ΔΔG for other inhibitors, FEP+ correctly predicted a very narrow range of ΔΔGs for this drug. This is reflected in the MUE and RMSE of 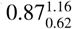 kcal/mol and 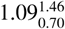 kcal/mol respectively, which are in-line with the MUEs and RMSEs for the other TKIs.

**Figure 4.**
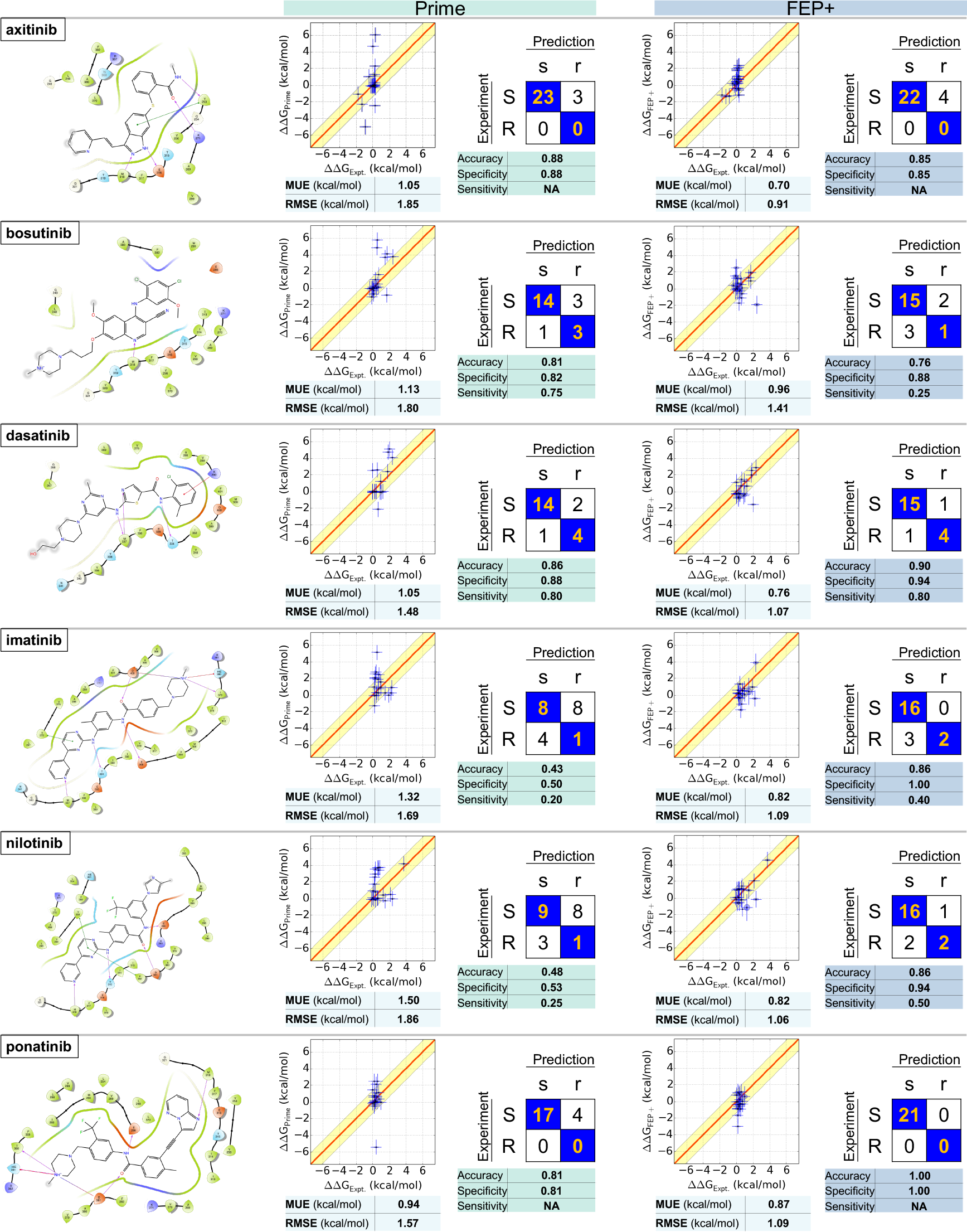
Physical modeling accuracy in computing the impact of clinical Abl mutations on selective inhibitor binding. Ligand interaction diagrams for six selective FDA-approved tyrosine kinase inhibitors (TKIs) for which co-crystal structures with Abl were available (left). Comparisons for clinically-observed mutations are shown for FEP+ (right) and Prime (left). For each ligand, computed *vs*. experimental binding free energies (ΔΔG)are plotted with MUE and RMSE (units of kcal/mol) depicted below. Truth tables are shown to the right. Rows denote *true* susceptible (S, ΔΔG < 1.36 kcal/mol) or resistant (R, ΔΔG > 1.36 kcal/mol) experimental classes using a 1.36 kcal/mol (10-fold change) threshold; columns denote *predicted* susceptible (s, ΔΔG ≤ 1.36 kcal/mol) or resistant (r, ΔΔG > 1.36 kcal/mol). Correct predictions populate diagonal elements (orange text), incorrect predictions populate off-diagonals. Accuracy, specificity, and sensitivity for two-class classification are shown below the truth table. Elliptical point sizes and error bars in the scatter plots depict estimated uncertainty/variability and error respectively (±σ) of FEP+ values (vertical size) and experimental values (horizontal size). Note: The sensitivity for axitinib and ponatinib is NA, because there is no resistant mutation for these two drugs.

### Understanding the origin of mispredictions

Resistance mutations that are mispredicted as susceptible (false negatives) are particularly critical because they might mislead the clinician or drug designer into believing the inhibitor will remain effective against the target. Which resistance mutations did FEP+ mispredict as susceptible? Nine mutations were classified by FEP+ to be susceptible when experimentally measured ΔpIC_50_ data indicate the mutations should have increased resistance according to our 10-fold affinity cutoff for resistance. Notably, the 95% confidence intervals for five of these mutations spanned the 1.36 kcal/mol threshold, indicating these misclassifications are not statistical significant when the experimental error and statistical uncertainty in FEP+ are accounted for: bosutinib/L248R 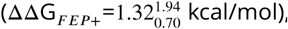 imatinib/E255K 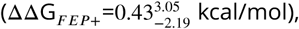 imatinib/Y253F 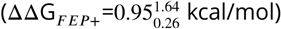, and nilotinib/Y253F 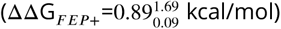. The bosutinib/V299L mutation was also not significant because the experimental 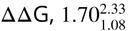 kcal/mol, included the 1.36 kcal/mol cutoff; the value of ΔΔG predicted by FEP+ for this mutation was 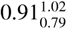 kcal/mol, the upper bound of the predicted value was within 0.06 kcal/mol of the lower bound of the experimental value.

Four mutations, however, were misclassified to a degree that is statistically significant given their 95% confidence intervals: dasatinib/T315A, bosutinib/T315I, imatinib/E255V, and nilotinib/E255V. For dasatinib/T315A, although the T315A mutations for bosutinib, imatinib, nilotinib, and ponatinib were correctly classified as susceptible, the predicted free energy changes for these four TKIs were consistently much more negative than the corresponding experimental measurements, just as for dasatinib/T315A, indicating there might be a generic driving force contributing to the errors in T315A mutations for these five TKIs. Abl is known to be able to adopt many different conformations (including DFG-in and DFG-out), and it is very likely that the T315A mutation will induce conformational changes in the apo protein [50], which was not adequately sampled in the relatively short simulations, leading to the errors for T315A mutations for these TKIs. By comparison, the T315I mutations for axitinib, bosutinib, imatinib, nilotinib, and ponatinib were all accurately predicted with the exception of bosutinib/T315I being the only misprediction, suggesting an issue specific to bosutinib. The complex electrostatic interactions between the 2,4-dichloro-5-methoxyphenyl ring in bosutinib and the adjacent positively charged amine of the catalytic Lys271 may not be accurately captured by the fixed-charge OPLS3 force field, leading to the misprediction for bosutinib/T315I mutation.

Insufficient sampling might also belie the imatinib/E255V and nilotinib/E255V mispredictions because they reside in the highly flexible P-loop. Since E255V was a charge change mutation, we utilized a workflow that included a transmutable explicit ion (see Methods). The distribution of these ions in the simulation box around the solute might not have converged to their equilibrium state on the relatively short timescale of our simulations (5 ns), and the insufficient sampling of ion distributions coupled with P-loop motions might lead to misprediction of these two mutations.

### How accurately can the impact of mutations be predicted for docked TKIs?

To assess the potential for utilizing physics-based approaches in the absence of a high-resolution experimental structure, we generated models of Abl bound to two TKIs—erlotinib and gefinitib—for which co-crystal structures with wild-type kinase are not currently available. In ***Figure 5***, we show the Abl:erlotinib and Abl:gefitinib complexes that were generated using a docking approach (Glide-SP, see Methods). These two structures were aligned against the co-crystal structures of EGFR:erlotinib and EGFR:gefinitib to highlight the structural similarities between the binding pockets of Abl and EGFR and the TKI binding mode in Abl versus EGFR. As an additional test of the sensitivity of FEP+ to system preparation, a second set of Abl:erlotinib and Abl:gefitinib complexes was generated in which crystallographic water coordinates were transferred to the docked inhibitor structures (see Methods).

Alchemical free-energy simulations were performed on 13 mutations between the two complexes; 7 mutations for erlotinib and 6 mutations for gefitinib. The quantitative accuracy of FEP+ in predicting the value of ΔΔG was excellent—MUE and RMSE of 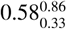 kcal/mol and 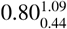 kcal/mol respectively if crystal waters are omitted, and 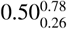 kcal/mol and 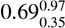 kcal/mol if crystal waters were restored after docking. Encouragingly, these results indicate that our initial models of Abl bound to erlotinib and gefitinib were reliable because the accuracy and dependability of our FEP+ calculations were not sensitive to crystallographic waters. Our secondary concern was the accuracy with which the approach classified mutations as resistant or susceptible.

While the results presented in (***Figure 5***) indicate that FEP+ is capable of achieving good quantitative accuracy when a co-crystal structure is unavailable, it is important to understand why a mutation was predicted to be susceptible but was determined experimentally to be resistant. F317I was the one mutation that increased resistance to erlotinib (or gefitinib) because it destabilized binding by more than 1.36 kcal/mol— 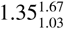 kcal/mol (gefitinib) and 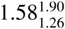 kcal/mol (erlotinib), but the magnitude of the experimental uncertainty means we are unable to confidently discern whether this mutation induces more than 10-fold resistance to either TKI. Therefore, the one misclassification by FEP+ in ***Figure 5*** is not statistically significant and the classification metrics presented there underestimate the nominal performance of this alchemical free-energy method.

**Figure 5.**
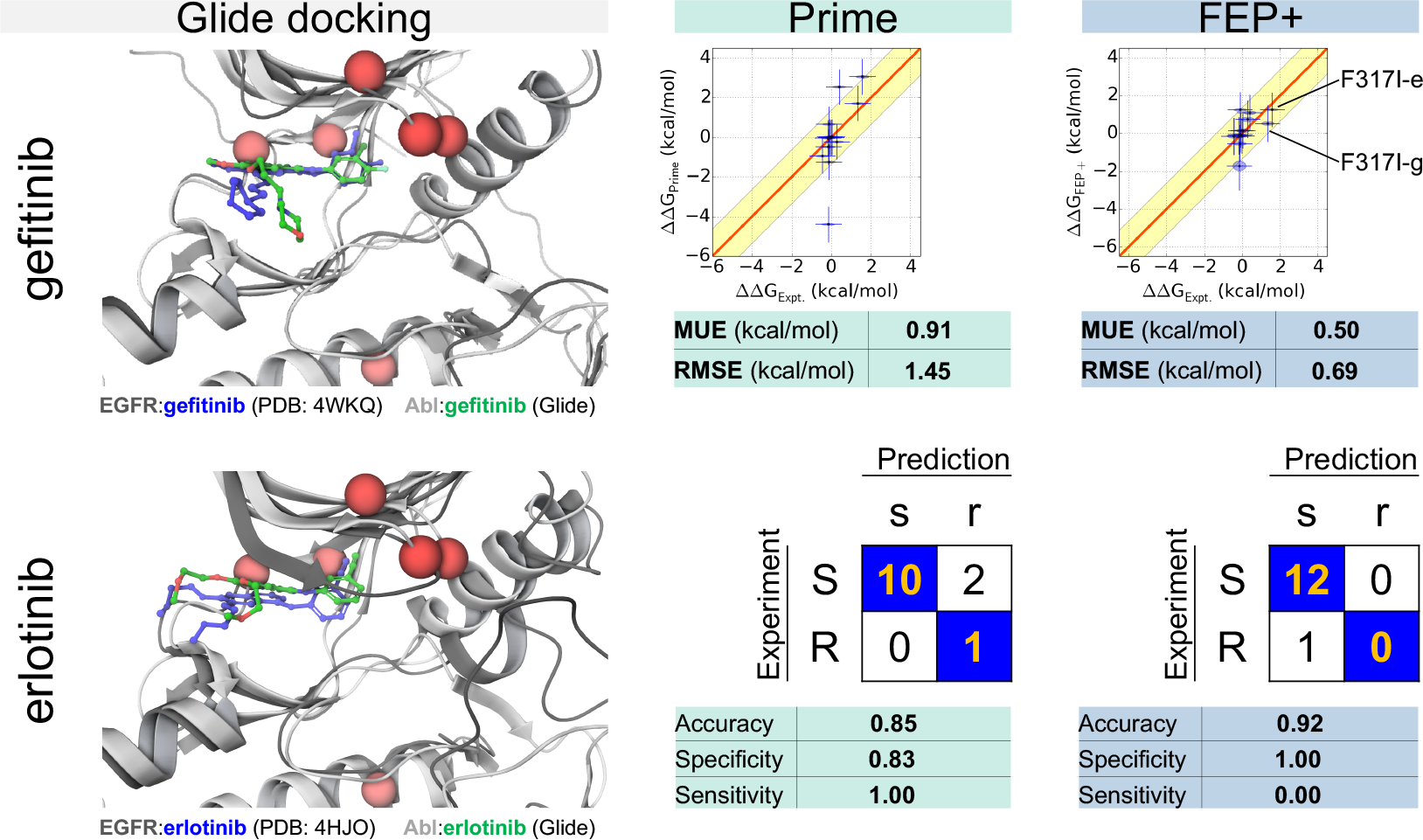
Predicting resistance mutations using FEP+ for inhibitors for which co-crystal structures with wild-type kinase are not available. The docked pose of Abl:erlotinib is superimposed on the co-crystal structure of EGFR:erlotinib; erlotinib docked to Abl (light gray) is depicted in green and erlotinib bound to EGFR (dark gray) is depicted in blue. The docked pose of Abl:gefitinib is superimposed on the co-crystal structure of EGFR:gefitinib; gefitinib docked to Abl (light gray) is depicted in green and gefitinib bound to EGFR (dark gray) is depicted in blue. The locations of clinical mutants for each inhibitor are highlighted (red spheres). The overall RMSEs and MUEs for Prime (center) and FEP+ (right) and two-class accuracies are also shown in the figure. Computed free energy changes due to the F317I mutation for erlotinib (-e) and gefitinib (-g) are highlighted in the scatter plot. FEP+ results are based on the docked models prepared with crystal waters added back while the Prime (an implicit solvent model) results are based on models without crystallographic water.

## Discussion

### Physics-based modeling can reliably predict when a mutation elicits resistance to therapy

The results presented in this work are summarized in ***Table 2***. The performance metrics summarized in ***Table 2*** indicates that the set of 131 mutations for the six TKIs in which co-crystal structures were available is on par with the complete set (144 mutations), which included results based on Abl:TKI complexes generated from docking models. The performance results for the 13 mutations for the two TKIs (erlotinib and gefitinib) in which co-crystal structures were unavailable exhibited good quantitative accuracy (MUE and RMSE) and good classification power.

**Table 2.** Summary of FEP+ and Prime statistics in predicting mutational resistance or sensitivity to FDA-approved TKIs.

| Dataset | Method | N<sub>quant</sub> | MUE<br>(kcal/mol) | RMSE<br>(kcal/mol) | N<sub>class</sub> | Accuracy | Specificity | Sensitivity |
| --- | --- | --- | --- | --- | --- | --- | --- | --- |
| all | FEP+ | 142 | 0.79<sup>0.92</sup><sub>0.67</sub> | 1.07<sup>1.26</sup><sub>0.89</sub> | 144 | 0.88<sup>0.93</sup><sub>0.82</sub> | 0.94<sup>0.98</sup><sub>0.89</sub> | 0.47<sup>0.69</sup><sub>0.25</sub> |
| all | Prime | 142 | 1.14<sup>1.35</sup><sub>0.93</sub> | 1.70<sup>1.98</sup><sub>1.40</sub> | 144 | 0.73<sup>0.80</sup><sub>0.66</sub> | 0.76<sup>0.84</sup><sub>0.69</sub> | 0.53<sup>0.76</sup><sub>0.30</sub> |
| xtals | FEP+ | 129 | 0.82<sup>0.95</sup><sub>0.69</sub> | 1.11<sup>1.30</sup><sub>0.91</sub> | 131 | 0.87<sup>0.92</sup><sub>0.81</sub> | 0.93<sup>0.97</sup><sub>0.88</sub> | 0.50<sup>0.74</sup><sub>0.29</sub> |
| xtals | Prime | 129 | 1.16<sup>1.37</sup><sub>0.96</sub> | 1.72<sup>2.00</sup><sub>1.41</sub> | 131 | 0.72<sup>0.79</sup><sub>0.64</sub> | 0.75<sup>0.83</sup><sub>0.67</sub> | 0.50<sup>0.73</sup><sub>0.25</sub> |
| axitinib | FEP+ | 26 | 0.70<sup>0.93</sup><sub>0.50</sub> | 0.91<sup>1.14</sup><sub>0.64</sub> | 26 | 0.85<sup>0.96</sup><sub>0.69</sub> | 0.85<sup>0.96</sup><sub>0.69</sub> | NA |
| axitinib | Prime | 26 | 1.05<sup>1.71</sup><sub>0.53</sub> | 1.85<sup>2.61</sup><sub>0.96</sub> | 26 | 0.88<sup>1.00</sup><sub>0.73</sub> | 0.88<sup>1.00</sup><sub>0.73</sub> | NA |
| bosutinib | FEP+ | 21 | 0.96<sup>1.42</sup><sub>0.55</sub> | 1.41<sup>1.97</sup><sub>0.77</sub> | 21 | 0.76<sup>0.95</sup><sub>0.57</sub> | 0.88<sup>1.00</sup><sub>0.71</sub> | 0.25<sup>1.00</sup><sub>0.00</sub> |
| bosutinib | Prime | 21 | 1.13<sup>1.83</sup><sub>0.60</sub> | 1.80<sup>2.62</sup><sub>0.92</sub> | 21 | 0.81<sup>0.95</sup><sub>0.62</sub> | 0.82<sup>1.00</sup><sub>0.62</sub> | 0.75<sup>1.00</sup><sub>0.00</sub> |
| dasatinib | FEP+ | 20 | 0.76<sup>1.13</sup><sub>0.49</sub> | 1.07<sup>1.57</sup><sub>0.59</sub> | 21 | 0.90<sup>1.00</sup><sub>0.76</sub> | 0.94<sup>1.00</sup><sub>0.79</sub> | 0.80<sup>1.00</sup><sub>0.33</sub> |
| dasatinib | Prime | 20 | 1.05<sup>1.54</sup><sub>0.61</sub> | 1.48<sup>1.92</sup><sub>0.95</sub> | 21 | 0.86<sup>1.00</sup><sub>0.71</sub> | 0.88<sup>1.00</sup><sub>0.69</sub> | 0.80<sup>1.00</sup><sub>0.33</sub> |
| imatinib | FEP+ | 20 | 0.82<sup>1.15</sup><sub>0.53</sub> | 1.09<sup>1.43</sup><sub>0.69</sub> | 21 | 0.86<sup>1.00</sup><sub>0.71</sub> | 1.00<sup>1.00</sup><sub>1.00</sub> | 0.40<sup>0.83</sup><sub>0.00</sub> |
| imatinib | Prime | 20 | 1.32<sup>1.81</sup><sub>0.91</sub> | 1.69<sup>2.26</sup><sub>1.15</sub> | 21 | 0.43<sup>0.67</sup><sub>0.24</sub> | 0.50<sup>0.75</sup><sub>0.25</sub> | 0.20<sup>0.67</sup><sub>0.00</sub> |
| nilotinib | FEP+ | 21 | 0.82<sup>1.12</sup><sub>0.57</sub> | 1.06<sup>1.39</sup><sub>0.69</sub> | 21 | 0.86<sup>1.00</sup><sub>0.67</sub> | 0.94<sup>1.00</sup><sub>0.80</sub> | 0.50<sup>1.00</sup><sub>0.00</sub> |
| nilotinib | Prime | 21 | 1.50<sup>1.97</sup><sub>1.06</sub> | 1.86<sup>2.25</sup><sub>1.43</sub> | 21 | 0.48<sup>0.67</sup><sub>0.24</sub> | 0.53<sup>0.75</sup><sub>0.29</sub> | 0.25<sup>1.00</sup><sub>0.00</sub> |
| ponatinib | FEP+ | 21 | 0.87<sup>1.16</sup><sub>0.62</sub> | 1.09<sup>1.46</sup><sub>0.70</sub> | 21 | 1.00<sup>1.00</sup><sub>1.00</sub> | 1.00<sup>1.00</sup><sub>1.00</sub> | NA |
| ponatinib | Prime | 21 | 0.94<sup>1.54</sup><sub>0.50</sub> | 1.57<sup>2.44</sup><sub>0.69</sub> | 21 | 0.81<sup>0.95</sup><sub>0.62</sub> | 0.81<sup>0.95</sup><sub>0.62</sub> | NA |
| Glide | FEP+ | 13 | 0.50<sup>0.78</sup><sub>0.26</sub> | 0.69<sup>0.97</sup><sub>0.35</sub> | 13 | 0.92<sup>1.00</sup><sub>0.77</sub> | 1.00<sup>1.00</sup><sub>1.00</sub> | 0.00<sup>0.00</sup><sub>0.00</sub> |
| Glide | Prime | 13 | 0.91<sup>1.56</sup><sub>0.39</sub> | 1.45<sup>2.22</sup><sub>0.54</sub> | 13 | 0.85<sup>1.00</sup><sub>0.62</sub> | 0.83<sup>1.00</sup><sub>0.58</sub> | 1.00<sup>1.00</sup><sub>0.00</sub> |

N<sub>quant</sub>: Number of mutations for which quantitative metrics were evaluated; N<sub>class</sub>: Number mutations for which classification metrics were evaluated; All: All mutations; xtals: All mutations for which co-crystal structures were available; Glide: erlotinib and gefitinib
Accuracy, specificity, and sensitivity were computed to assess two-class prediction performance:
*resistant* ( $\Delta\Delta G > 1.36$ kcal/mol) or *susceptible* ( $\Delta\Delta G \leq 1.36$ kcal/mol).
95% CIs (sub-/superscripts) were estimated from 1000 bootstrap replicates. Note: The sensitivity for axitinib and ponatinib is NA, because there is no resistant mutation for these two drugs.

Overall (N=144), the MM-GBSA approach Prime classified mutations with good accuracy 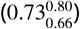 and specificity 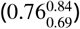 while the alchemical approach FEP+ was a significant improvement in classification accuracy 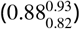 and specificity 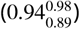. The quantitative accuracy with which Prime was able to predict the experimentally measured change in Abl:TKI binding (N=142) characterized by RMSE and MUE was 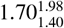 kcal/mol and 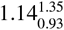 kcal/mol respectively. In stark contrast, the quantitative accuracy of FEP+ was statistically superior to Prime with an RMSE and an MUE of 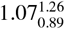 kcal/mol and 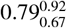 kcal/mol respectively.

From the perspective of a clinician, classification rate would be an important metric to measure the predictive power of technologies such as Prime and FEP+. To test the hypothesis that reducing the large spread in Prime predictions could improve its classification rate, we scaled the computed relative free energies (by 1/2, 1/3, and by 0.23, which was the optimal factor that gives lowest RMSE) and recalculated the classification metrics (***Table S8***). As expected, the MUE and RMSE were improved but the specificity of Prime was drastically diminished; as MUE and RMSE improved, it became increasingly unable to identify resistance mutations. Scaling FEP+ eliminated its sensitivity and a naïve model (where all free energies were set to 0.00 kcal/mol) had zero sensitivity. Lastly, we constructed a consensus model in which free energies were a weighted average of scaled Prime and FEP+. However, this model also had no sensitivity. It appears difficult to improve upon the predictive power of FEP+ by statistical operations.

To address the impact of picking a cutoff to classify predicted free energies as resistant or sensitizing, we computed ROC curves for the various predicted datasets: Prime (scaled and non-scaled), FEP+ (scaled and non-scaled), naïve model, and consensus model (constructed from scaled Prime and scaled FEP+, see above). ROC curves are independent of a linear transformation on the predicted dataset. Therefore, ROC curves and ROC-AUCs for scaled and non-scaled Prime were identical, as well as scaled and non-scaled FEP+. ROC curves for these six sets of predictions are presented in Supplementary ***Figure S3***. ROC-AUC for FEP+ was 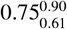 (n=144); ROC-AUC for Prime was 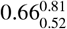 (n=144); ROC-AUCs for the naïve model and consensus model were 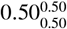 (n=144) and 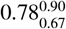 (n=144) respectively. These results show that Prime apparently has poor discriminatory power (ROC-AUC in [0.6,0.7]) while FEP+ apparently has fair discriminatory power (ROC-AUC in [0.7,0.8]).

### Hierarchical Bayesian model estimates global performance (N=144)

A hierarchical Bayesian approach was developed to estimate the intrinsic accuracy of the models when the noise in the experimental and predicted values of ΔΔG was accounted for. Utilizing this approach, the MUE and RMSE for Prime was found to be 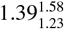 kcal/mol and 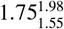 kcal/mol (N=142) respectively. The accuracy, specificity, and sensitivity of Prime was found using this method to be 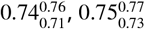, and 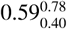 (N=144) respectively. The MUE and RMSE of FEP+ was found to be 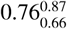 kcal/mol and 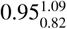 kcal/mol (N=142) respectively, which is significantly better than Prime. Likewise, a clearer picture of the true classification accuracy, specificity, and sensitivity of FEP+ was found—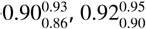, and 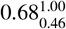 respectively.

### Examining the physical and chemical features of outliers

#### Current alchemical approaches neglect effects that will continue to improve accuracy

The high accuracy of FEP+ is very encouraging, and the accuracy can be further improved with more accurate modeling of a number of physical chemical effects not currently considered by the method. While highly optimized, the fixed-charged OPLS3 [25] force field can be further improved by explicit consideration of polarizability effects [51], as hinted by some small-scale benchmarks [52]. These features could be especially important for bosutinib, whose 2,4-dichloro-5-methoxyphenyl ring is adjacent to the positively charged amine of the catalytic Lys271. Many simulation programs also utilize a long-range isotropic analytical dispersion correction intended to correct for the truncation of dispersion interactions at finite cutoff, which can induce an error in protein-ligand binding free energies that depends on the number of ligand heavy atoms being modified [53]; recently, efficient Lennard-Jones PME methods [54, 55] and perturbation schemes [53] have been developed that can eliminate the errors associated with this truncation. While the currently employed methodology for alchemical transformations involving a change in system charge (see Methods) reduces artifacts that depend on the simulation box size and periodic boundary conditions, the explicit ions that were included in these simulations may not have sufficiently converged to their equilibrium distributions in these relatively short simulations. Kinases and their inhibitors are known to possess multiple titratable sites with either intrinsic or effective p*K*_*a*_s near physiological pH, while the simulations here treat protonation states and proton tautomers fixed throughout the bound and unbound states; the accuracy of the model can be further improved with the protonation states or tautomers shift upon binding or mutation considered [56,57]. Similarly, some systems display significant salt concentration dependence [58], while the simulations for some systems reported here did not rigorously mimic all aspects of the experimental conditions of the cell viability assays.

#### Experimentally observed IC_50_ changes can be caused by other physical mechanisms

While we have shown that predicting the direct impact of mutations on the binding affinity of ATP-competitive tyrosine kinase inhibitors for a single kinase conformation has useful predictive capacity, many additional physical effects that can contribute to cell viability are not currently captured by examining only the predicted change in inhibitor binding affinity. For example, kinase missense mutations can also shift the populations of kinase conformations (which may affect ATP and inhibitor affinities differentially), modulate ATP affinity, modulate affinity for protein substrate, or modulate the ability of the kinase to be regulated or bounded by scaffolding proteins. These physical mechanisms might affect the IC_50_s of cell viability assays but not necessarily the binding affinity of the inhibitors. While many of these effects are in principle tractable by physical modeling in general (and alchemical free energy methods in particular), it is valuable to examine our mispredictions and outliers to identify whether any of these cases is likely to induce resistance (as observed by ΔpIC_50_ shifts) by one of these alternative mechanisms.

#### Other physical mechanisms of resistance are likely similarly computable

A simple threshold of 10-fold TKI affinity change is a crude metric for classifying resistance or susceptibility due to the myriad biological factors that contribute to the efficacy of a drug in a person. Except for affecting the binding affinity of inhibitors, missense mutations can also cause drug resistance through other physical mechanisms including induction of splice variants or alleviation of feedback. While the current study only focused on the effect of mutation on drug binding affinity, resistance from these other physical mechanisms could be similarly computed using physical modeling. For example, some mutations are known to activate the kinase by increasing affinity to ATP, which could be computed using the same thermodynamic cycle utilized here for inhibitors.

## Conclusion

Revolutionary changes in computing power—especially the arrival of inexpensive graphics processors (GPUs)—and software automation have enabled alchemical free-energy calculations to impact drug discovery and life sciences projects in previously unforeseen ways. In this communication, we tested the hypothesis that FEP+, a fully-automated relative-alchemical free-energy workflow, had reached the point where it can accurately and reliably predict how clinically-observed mutations in Abl kinase alter the binding affinity of eight FDA-approved TKIs. To establish the potential predictive impact of current-generation alchemical free energy calculations—which incorporate entropic and enthalpic effects and the discrete nature of aqueous solvation—compared to a simpler physics-based approach that also uses modern forcefields but scores a single minimized conformation, we employed a second physics-based approach (Prime). This simpler physics-based model, which uses an implicit model of solvation to score the energetic changes in interaction energy that arise from the mutation, was able to capture a useful amount of information to achieve substantial predictiveness with an MUE of 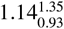 kcal/mol (N=142), RMSE of 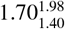 kcal/mol respectively (N=142), and classification,cation accuracy of 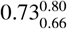 (N=144). Surpassing these good results we went on to demonstrate that FEP+ is able to achieve superior predictive performance—MUE of 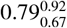 kcal/mol (N=142), RMSE of 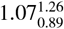 kcal/mol (N=142), and classification accuracy of 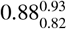 (N=144). While future enhancements to the workflows for Prime and FEP+ to account for additional physical and chemical effects are likely to improve predictive performance further, the present results are of sufficient quality and achievable on a sufficiently rapid timescale (with turnaround times **~**6 hours/calculation) to impact research projects in drug discovery and the life sciences. With exponential improvements in computing power, we anticipate the domains of applicability for alchemical free-energy methods such as FEP+ will take on increasingly integrated roles to impact projects. This work illustrates how the domain of applicability for alchemical free-energy methods is much larger than previously appreciated, and might further be found to include new areas as research progresses: aiding clinical decision-making in the selection of first- or second-line therapeutics guided by knowledge of likely subclonal resistance; identifying other selective kinase inhibitors (or combination therapies) to which the mutant kinase is susceptible; supporting the selection of candidate molecules to advance to clinical trials based on anticipated activity against likely mutations; facilitating the enrollments of patients in mechanism-based basket trials; and generally augmenting the armamentarium of precision oncology.

## Methods

### System preparation

All system preparation utilized the Maestro Suite (Schrödinger) version 2016-4. Comparative modeling to add missing residues using a homologous template made use of the Splicer tool, while missing loops modeled without a template used Prime. All tools employed default settings unless otherwise noted. The Abl wild-type sequence used in building all Abl kinase domain models utilized the ABL1_HUMAN Isoform IA (P00519-1) UniProt gene sequence spanning S229–K512. Models were prepared in non-phosphorylated form. We used a residue indexing convention that places the Thr gatekeeper residue at position 315 to match common usage; an alternate indexing convention utilized in experimental X-ray structures for Abl:imatinib (PDB: 1OPJ) [59] and Abl:dasatinib (PDB: 4XEY) [60] was adjusted to match our convention.

#### Complexes with co-crystal structures

Chain B of the experimental structure of Abl:axitinib (PDB: 4WA9) [44] was used, and four missing residues at the N-and C-termini were added using homology modeling with PDB 3IK3 [61] as the template following alignment of the respective termini of the kinase domain. Chain B was selected because chain A was missing an additional 3 and 4 residues at the N-and C-termini, respectively, in addition to 3- and 20-residue loops, both of which were resolved in chain B. All missing side chains were added with Prime. The co-crystal structure of Abl:bosutinib (PDB: 3UE4) [62] was missing 4 and 10 N-and C-terminal residues respectively in chain A that were built using homology modeling with 3IK3 as the template. All loops were resolved in chain A (chain B was missing two residues in the P-loop, Q252 and Y253). All missing side chains were added with Prime. The co-crystal structure of Abl:dasatinib (PDB: 4XEY) [60] was missing 2 and 9 N-and C-terminal residues, respectively, that were built via homology modeling using 3IK3 as the template. A 3 residue loop was absent in chain B but present in chain A; chain A was chosen. The co-crystal structure of Abl:imatinib (PDB: 1OPJ) [59] had no missing loops. Chain B was used because chain A was missing two C-terminal residues that were resolved in chain B. A serine was present at position 336 (index 355 in the PDB file) and was mutated to asparagine using Prime to match the human wild-type reference sequence (P00519-1). The co-crystal structure of Abl:nilotinib (PDB: 3CS9) [63] contained four chains in the asymmetric unit all of which were missing at least one loop. Chain A was selected because its one missing loop involved the fewest number of residues of the four chains; chain A was missing 4 and 12 N-and C-terminal residues, respectively, that were built using homology modeling with 3IK3 as the template. A 4-residue loop was missing in chain A (chain B and C were missing two loops, chain D was missing a five residue loop) that was built using Prime. The co-crystal structure of Abl:ponatinib (PDB: 3OXZ) [64] contained only one chain in the asymmetric unit. It had two missing loops, one 4 residues (built using Prime) and one 12 residues (built using homology modeling with 3OY3 [64] as the template). Serine was present at position 336 and was mutated to Asn using Prime to match the human wild-type reference sequence (P00519-1). Once the residue composition of the six Abl:TKI complexes were normalized to have the same sequence, the models were prepared using Protein Preparation Wizard. Bond orders were assigned using the Chemical Components Dictionary and hydrogen atoms were added. Missing side chain atoms were built using Prime. Termini were capped with N-acetyl (N-terminus) and N-methyl amide (C-terminus). If present, crystallographic water molecules were retained. Residue protonation states (e.g. Asp381 and Asp421) were determined using PROPKA[65] with a pH range of 5.0-9.0. Ligand protonation state was assigned using PROPKA with pH equal to the experimental assay. Hydrogen bonds were assigned by sampling the orientation of crystallographic water, Asn and Gln flips, and His protonation state. The positions of hydrogen atoms were minimized while constraining heavy atoms coordinates. Finally, restrained minimization of all atoms was performed in which a harmonic positional restraint (25.0 kcal/mol/Å2) was applied only to heavy atoms. ***Table S9*** summarizes the composition of the final models used for FEP.

#### Complexes without co-crystal structures

Co-crystal structures of Abl bound to erlotinib or gefitinib were not publicly available. To generate models of these complexes, Glide-SP [66] was utilized to dock these two compounds into an Abl receptor structure. Co-crystal structures of these two compounds bound to EGFR were publicly available and this information was used to obtain initial ligand geometries and to establish a reference binding mode against which our docking results could be structurally scored. The Abl receptor structure bound to bosutinib was used for docking because its structure was structurally similar to that of EGFR in the erlotinib-(PDB: 4HJO) [67] and gefitinib-bound (PDB: 4WKQ) [68] co-crystal structures. Abl was prepared for docking by using the Protein Preparation Wizard (PPW) with default parameters. Crystallographic waters were removed but their coordinates retained for a subsequent step in which they were optionally reintroduced. Erlotinib and gefitinib protonation states at pH 7.0±2.0 were determined using Epik [69]. Docking was performed using the Glide-SP workflow. The receptor grid was centered on bosutinib. The backbone NH of Met318 was chosen to participate in a hydrogen bonding constraint with any hydrogen bond donor on the ligand. The hydroxyl of T315 was allowed to rotate in an otherwise rigid receptor. Ligand docking was performed with enhanced sampling; otherwise default settings were used. Epik state penalties were included in the scoring. The 16 highest ranked (Glide-SP score) poses were retained for subsequent scoring. To determine the docked pose that would be subsequently used for free energy calculations, the ligand heavy-atom RMSD between the 16 poses and the EGFR co-crystal structures (PDB IDs 4HJO and 4WKQ) was determined. The pose in which erlotinib or gefitinib most structurally resembled the EGFR co-crystal structure (lowest heavy-atom RMSD) was chosen as the pose for subsequent FEP+. Two sets of complex structures were subjected to free energy calculations to determine the effect of crystal waters: In the first set, without crystallographic waters, the complexes were prepared using Protein Prep Wizard as above. In the second set, the crystallographic waters removed prior to docking were added back, and waters in the binding pocket that clashed with the ligand were removed.

### Force field parameter assignment

The OPLS3 forcefield [25] version that shipped with Schrödinger Suite release 2016-4 was used to parameterize the protein and ligand. Torsion parameter coverage was checked for all ligand fragments using Force Field Builder. The two ligands that contained a fragment with a torsion parameter not covered by OPLS3 were axitinib and bosutinib; Force Field Builder was used to obtain these parameters. SPC parameters [70] were used for water. For mutations that change the net change of the system, counterions were included to neutralize the system with additional Na+ and Cl- ions added to achieve 0.15 M excess to mimic the solution conditions of the experimental assay.

### Prime (MM-GBSA)

Prime was used to predict the geometry of mutant side chains and to calculate relative changes in free energy using MM-GBSA single-point estimates [39]. VSGB [71] was used as the implicit solvent model to calculate the solvation free energies for the four states (complex/wild-type, complex/mutant, apo protein/wild-type, and apo protein/mutant) and ΔΔG calculated using the thermodynamic cycle depicted in ***Figure 1***b. Unlike FEP (see below), which simulates the horizontal legs of the thermodynamic cycle, MM-GBSA models the vertical legs by computing the interaction energy between the ligand and protein in both wild-type and mutant states, subtracting these to obtain the ΔΔG of mutation on the binding free energy.

### Alchemical free energy perturbation calculations using FEP+

Alchemical free energy calculations were performed using the FEP+ tool in the Schrödinger Suite version 2016-4, which offers a fully automated workflow requiring only an input structure (wild-type complex) and specification of the desired mutation. The default protocol was used throughout: It assigns protein and ligand force field parameters (as above), generates a dual-topology [72] alchemical system for transforming wild-type into mutant protein (whose initial structure is modeled using Prime), generates the solvent-leg endpoints (wild-type and mutant apo protein), and constructs intermediate windows spanning wild-type and mutant states. Simulations of the apo protein were setup by removing the ligand from the prepared complex (see System Preparation) followed by an identical simulation protocol as that used for the complex. Charge-conserving mutations utilized 12 *λ* windows (24 systems) while charge-changing mutations utilized 24 *λ* windows (48 systems). Each system was solvated in an orthogonal box of explicit solvent (SPC water [70]) with box size determined to ensure that solute atoms were no less than 5 Å (complex leg) or 10 Å (solvent leg) from an edge of the box. For mutations that change the net charge of the system, counterions were included to neutralize the charge of the system, and additional Na+ and Cl- ions added to achieve 0.15 M excess NaCl to mimic the solution conditions of the experimental assay. The artifact in electrostatic interactions for charge change perturbations due to periodic boundary conditions in MD simulations are corrected based on the method proposed by Rocklin *et al.* [73].

System equilibration was automated. It followed the default 5-stage Desmond protocol: (i) 100 ps with 1 fs time steps of Brownian dynamics with positional restraints of solute heavy atoms to their initial geometry using a restraint force constant of 50 kcal/mol/Å^2^; this Brownian dynamics integrator corresponds to a Langevin integrator in the limit when τ → 0, modified to stabilize equilibration of starting configurations with high potential energies; particle and piston velocities were clipped so that particle displacements were limited to 0.1 Å, in any direction. (ii) 12 ps MD simulations with 1 fs time step using Langevin thermostat at 10 K with constant volume, using the same restraints; (iii) 12 ps MD simulations with 1 fs time step using Langevin thermostat and barostat [74] at 10 K and constant pressure of 1 atmosphere, using the same restraints; (iv) 12 ps MD simulations with 1 fs time step using Langevin thermostat and barostat at 300 K and constant pressure of 1 atmosphere, using the same restraints; (v) a final unrestrained equilibration MD simulation of 240 ps with 2 fs time step using Langevin thermostat and barostat at 300 K and constant pressure of 1 atmosphere. Electrostatic interactions were computed with particle-mesh Ewald (PME) [75] and a 9 Å cutoff distance was used for van de Waals interactions. The production MD simulation was performed in the NPT ensemble using the MTK method [76] with integration time steps of 4 fs, 4 fs, and 8 fs respectively for the bonded, near, and far interactions following the RESPA method [77] through hydrogen mass repartitioning [78]. Production FEP+ calculations utilized Hamiltonian replica exchange with solute tempering (REST) [79], with automated definition of the REST region. Dynamics were performed with constant pressure of 1 atmosphere and constant temperature of 300 K for 5 ns in which exchanges between windows was attempted every 1.2 ps.

Because cycle closure could not be used to reduce statistical errors via path redundancy [79], we instead performed mutational free energy calculations in triplicate by initializing dynamics with different random seeds. The relative free energies for each mutation in each independent run were calculated using BAR [80, 81] The reported ΔΔG was computed as the mean of the computed ΔΔG from three independent simulations. Triplicate simulations were performed in parallel using four NIVIDA Pascal Architecture GPUs per alchemical free-energy simulation (12 GPUs in total), requiring ~6 hours in total to compute ΔΔG.

### Obtaining ΔΔG from ΔpIC_50_ benchmark set data

Reference relative free energies were obtained from three publicly available sources of ΔpIC_50_ data (***Table 1***). Under the assumption of Michaelis-Menten binding kinetics (pseudo first-order, but relative free energies are likely consistent), the inhibitor is competitive with ATP (***Equation 1***). This assumption has been successfully used to estimate relative free energies [37, 82–84] using the relationship between IC_50_ and competitive inhibitor affinity *K*_*i*_,

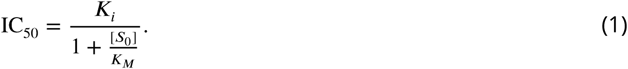

If the Michaelis constant for ATP (*K*_*M*_) is much larger than the initial ATP concentration *S*_0_, the relation in ***Equation 1*** will tend towards the equality IC_50_ = *K*_i_. The relative change in binding free energy of Abl:TKI binding due to protein mutation is simply,

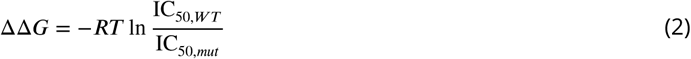

where IC_50,*WT*_ is the IC_50_ value for the TKI binding to the wild-type protein and IC_50,*mut*_ is the IC_50_ value for the mutant protein. *R* is the ideal gas constant and *T* is taken to be room temperature (300 K).

As alluded to above, relating ΔpIC_50_s to ΔΔGs assumes that the Michaelis constant for ATP is much larger than the initial concentration of ATP, and that the experimentally observed ΔpIC_50_ change is solely from changes in kinase:TKI binding affinity. In practice, not all of these assumptions may hold. For example, the experimentally observed ΔpIC_50_ might depend on the metabolism of drugs, and for drugs with different mechanisms of action than directly binding to the kinase binding pocket (e.g., binding to the transition structures of kinases, target gene amplification, up-/down-regulation of positive-/negative-feedback effectors, diminished synergism of pro-apoptotic machinery, decoupling of the target from cell survival circuits) [85, 86], their inhibition ability might not correlate well with binding affinity. However, the comparison between ΔpIC_50_ and Δ*K_D_* is presented in ***Figure 2***d, and this comparison indicates the assumptions we used to relate ΔpIC_50_ to ΔΔG are reasonable for the dataset we studied.

#### Assessing prediction performance

##### Quantitative accuracy metrics

Mean unsigned error (MUE) was calculated by taking the average absolute difference between predicted and experimental estimates of ΔΔG. Root-mean square error (RMSE) was calculated by taking the square root of the average squared difference between predicted and experimental estimates of ΔΔG. MUE depends linearly on errors such that large and small errors contribute equally to the average value, while RMSE depends quadratically on errors, magnifying their effect on the average value.

##### Truth tables

Two-class truth tables were constructed to characterize the ability of Prime and FEP+ to correctly classify mutations as susceptible (ΔΔG ≤ 1.36 kcal/mol) or resistant (ΔΔG > 1.36 kcal/mol), where the 1.36 kcal/mol threshold represents a 10-fold change in affinity. Accuracy was calculated as the fraction of all predictions that were correctly classified as sensitizing, neutral, or resistant. Sensitivity and specificity were calculated using a binary classification of resistant (ΔΔG > 1.36 kcal/mol) or susceptible (ΔΔG ≤ 1.36 kcal/mol). Specificity was calculated as the fraction of correctly predicted non-resistant mutations out of all truly susceptible mutations **S**. Sensitivity was calculated as the fraction of correctly predicted resistant mutations out of all truly resistant mutations, **R**. The number of susceptible mutations was 113 for axitinib, bosutinib, dasatinib, imatinib, nilotinib and ponatinib, and 12 for erlotinib and gefitinib; the number of resistant mutations **R** was 18 for axitinib, bosutinib, dasatinib, imatinib, nilotinib, and ponatinib, and 1 for erlotinib and gefitinib.

##### Consensus model

First, Prime and FEP+ (n=142) were scaled by minimizing their RMSE to experiment by optimizing slope using linear regression. The resulting (minimum) RMSE was used in a subsequent step to combine the scaled FEP+ and scaled Prime free energies with inverse-variance weighted averaging.

##### ROC

A ROC curve was generated by computing the true positive rate (sensitivity) and the true negative rate (specificity) when the classification cutoff differentiating resistant from sensitizing mutations is changed for (only) the predicted values of ΔΔG. Cutoffs were chosen by taking the minimum and maximum value of ΔΔG for a data set (Prime or FEP+), and iteratively computing specificity and sensitivity in steps of 0.001 kcal/mol, which by this definition will be in the range [0,1]. Experimental positives and negatives were classified with the 1.36 kcal/mol cutoff. ROC-AUC was computed using the trapezoidal rule.

##### Estimating uncertainties of physical-modeling results

95% symmetric confidence intervals (CI, 95%) for all performance metrics were calculated using bootstrap by resampling all datasets with replacement, with 1000 resampling events. Confidence intervals were estimated for all performance metrics and reported as 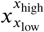 where *x* is the mean statistic calculated from the complete dataset (e.g. RMSE), and *x*_low_ and *x*_high_ are the values of the statistic at the 2.5^*th*^ and 97.5^*th*^ percentiles of the value-sorted list of the bootstrap samples. Uncertainty for ΔΔGs was computed by the standard deviation between three independent runs (using different random seeds to set initial velocities), where the 95% CI was [ΔΔG-1.96×σ_FEP+_, ΔΔG+1.96×σ_FEP+_] kcal/mol. 1σ used in plots for FEP+ and experiment; 0σ for Prime.

##### Bayesian hierarchical model to estimate intrinsic error

We used Bayesian inference to estimate the true underlying prediction error of Prime and FEP+ by making use of known properties of the experimental variability (characterized in ***Figure 2***) and statistical uncertainty estimates generated by our calculations under weak assumptions about the character of the error.

We presume the true free energy differences of mutation *i*, 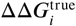 true, comes from a normal background distribution of unknown mean and variance,

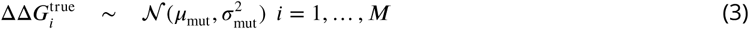

where there are *M* mutations in our dataset. We assign weak priors to the mean and variance

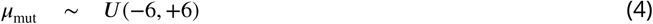

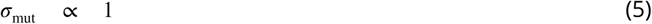

where we limit σ > 0.

We presume the true computational predictions (absent statistical error) differ from the (unknown) true free energy difference of mutation 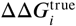 by normally-distributed errors with zero bias but standard deviation equal to the RMSE for either Prime or FEP+, the quantity we are focused on estimating:

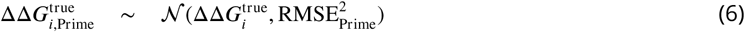

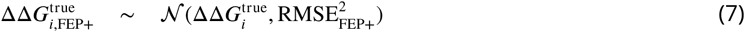

In the case of Prime, since the computation is deterministic, we actually calculate 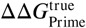 for each mutant. For FEP+, however, the computed free energy changes are corrupted by statistical error, which we also presume to be normally distributed with standard deviation σ_calc,i_

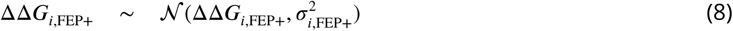

where ΔΔ*G*_i,FEP+_ is the free energy computed for mutant, *i* by FEP+, and σ_*i*,FEP+_ is the corresponding statistical error estimate.

The experimental data we observe is also corrupted by error, which we presume to be normally distributed with standard deviation σ_exp_:

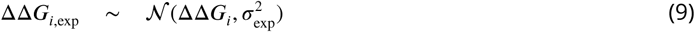

Here, we used an estimate of *K*_*d*_- and IC_50_-derived ΔΔ*G* variation derived from the empirical RMSE of 0.81 kcal/mol, where we took σ_exp_ ≈ 0.81 /√2 = 0.57 kcal/mol to ensure the difference between two random measurements of the same mutant would have an empirical RMSE of 0.81 kcal/mol.

Under the assumption that the true ΔΔ*G* is normally distributed and the calculated value differs from the true value via a normal error model, it can easily be shown that the MUE is related to the RMSE via

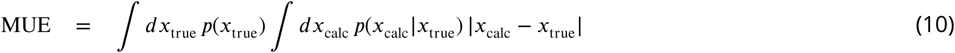

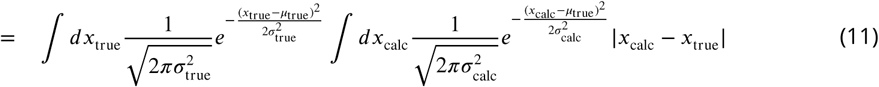

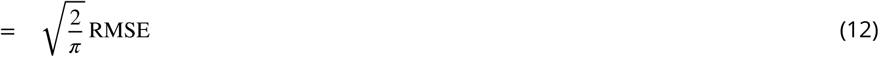

The model was implemented using PyMC3 [87], observable quantities were set to their computed or experimental values, and 500 samples drawn from the posterior (after discarding an initial 500 samples to burn-in) using the default NUTS sampler. Expectations and posterior predictive intervals were computed from the marginal distributions obtained from the resulting traces.

#### Data availability

Compiled experimental datasets, input files for Prime and FEP+ and computational results can be found at the following URL: https://goo.gl/6cC8Bu

#### Code availability

Scripts used for statistics analysis (including the Bayesian inference model) can be found at the following URL: https://goo.gl/6cC8Bu

## Acknowledgments

We thank Daniel Robinson (Schrödinger), Sonya M. Hanson (MSKCC), and Gregory A. Ross (MSKCC) for helpful discussions. JDC acknowledges support from NIH National Cancer Institute Cancer Center Core Grant P30 CA008748;JDC and SKA acknowledge support from the Sloan Kettering Institute, Cycle for Survival, and NIH grant R01 GM121505. KH acknowledges help from Wei Chen (Schrödinger) and Anthony Clark (Schrödinger) for instructions on running mutations changing the net charge of the system, and Simon Gao (Schrödinger) for assistance in computational resources.

## Author Contributions

KH, JDC, CN, RA, and LW designed the research; KH, SA, TS, and LW identified experimental datasets; KH and LW performed the simulations; KH, CN, SKA, SR, TS, RA, JDC, and LW analyzed the data; KH, JDC, SKA, and LW wrote the paper.

## Supplementary Information

**Figure S1.**
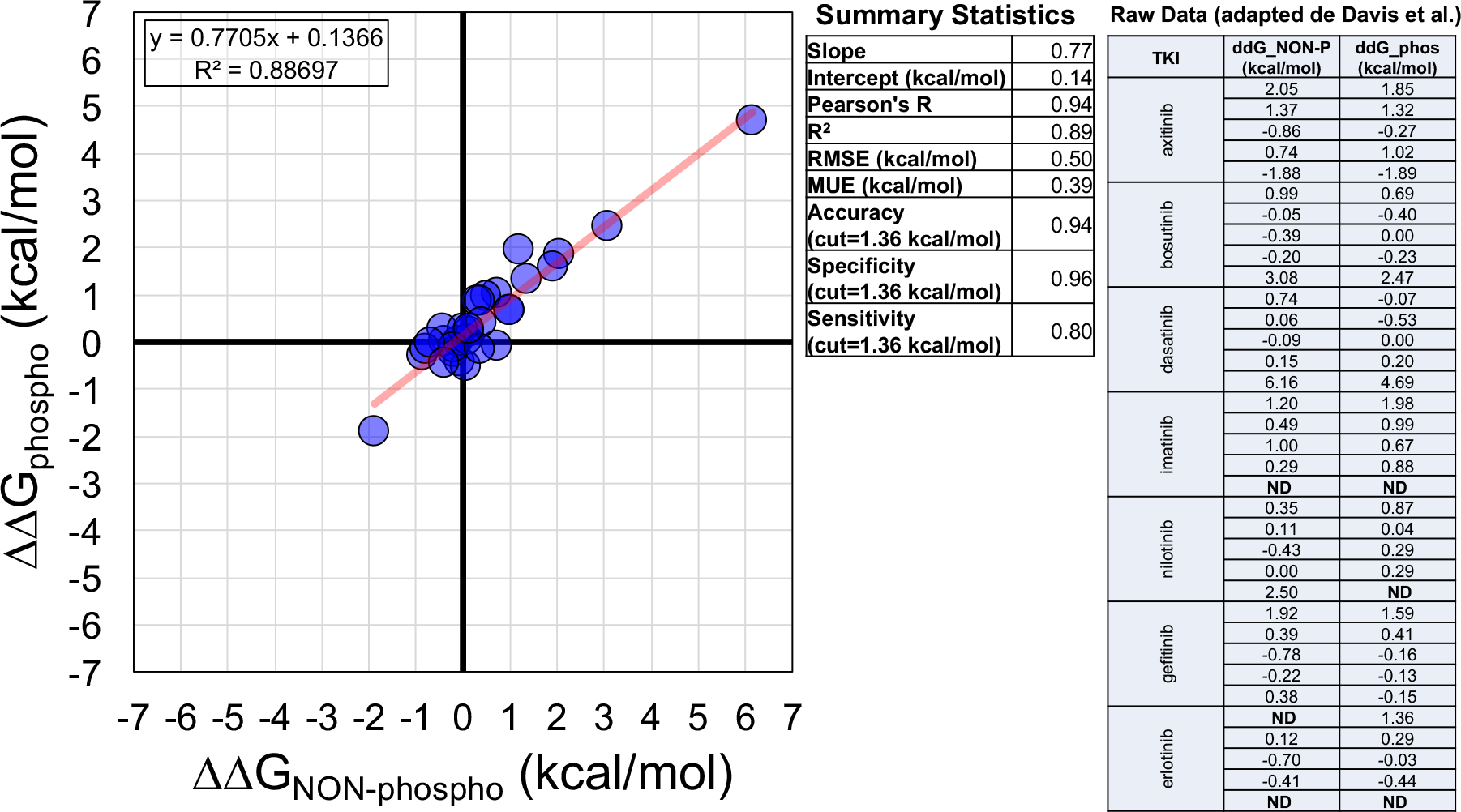
Comparison of 31 mutations for which phosphorylated and non-phosphorylated Δ*K_d_*s were available. Scatter plot compares ΔΔGs (derived from the Δ*K_d_*s) and contains the best-1t line with slope 0.77 and intercept 0.14. Summary statistics for this comparison are also shown. The raw ΔΔGs used for this comparison were adapted from [46]; kino-bead data for ponatinib was not available.

**Figure S2.**
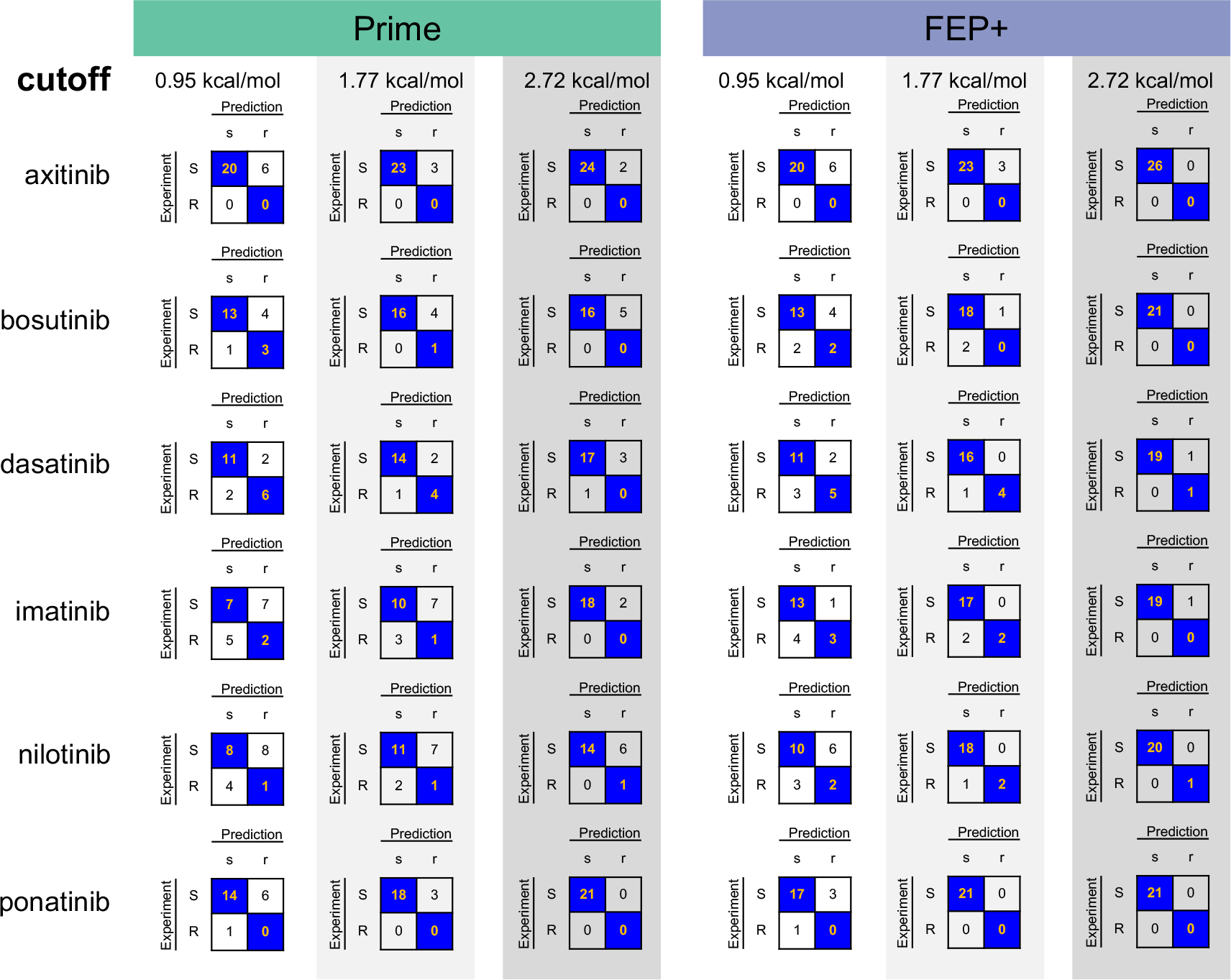
TKI-by-TKI truth tables with increasingly large classification cutoffs. Truth tables for the six TKIs (axitinib, bosutinib, dasatinib, imatinib, nilotinib, and ponatinib) using Prime (left, green) and FEP+ (right, blue) with classification cutoff values defining whether mutations are susceptible (S, experiment; s, prediction) or resistant (R, experiment; r, prediction). A mutation is susceptible if ΔΔG ≤ **cutoff** or resistant if ΔΔG > **cutoff**.

**Figure S3.**
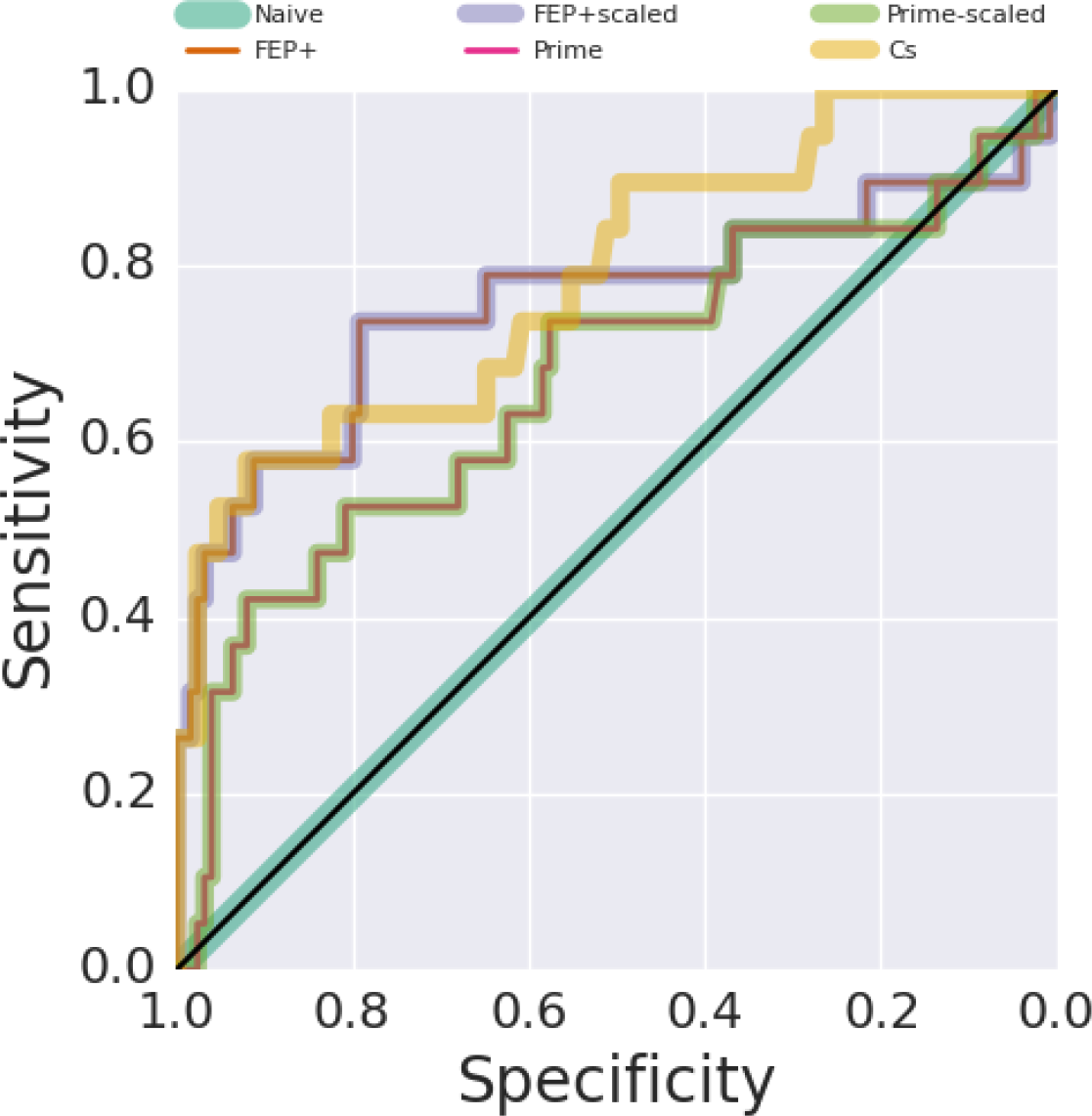
ROC curves for non-scaled and scaled FEP+, non-scaled and scaled Prime, a consensus model and a naïve model. ROC-AUC for scaled and non-scaled FEP+ was 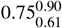 (n=144); ROC-AUC for scaled and non-scaled Prime was 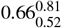 (n=144); ROC-AUCs for the naïve model and consensus model were 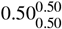 (n=144) and 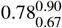 (n=144) respectively. Optimal scaling factors (a=0.34 for FEP+; a=0.23 for Prime) obtained using linear regression (m=142) were applied to the full dataset (n=144), which was used in this ROC analysis. ROC-AUC interpretations: [0.50,0.60], failure; [0.60,0.70], poor; [0.70,0.80], fair; [0.80,0.90], good; [0.90,1.00], excellent.

**Table S1.**
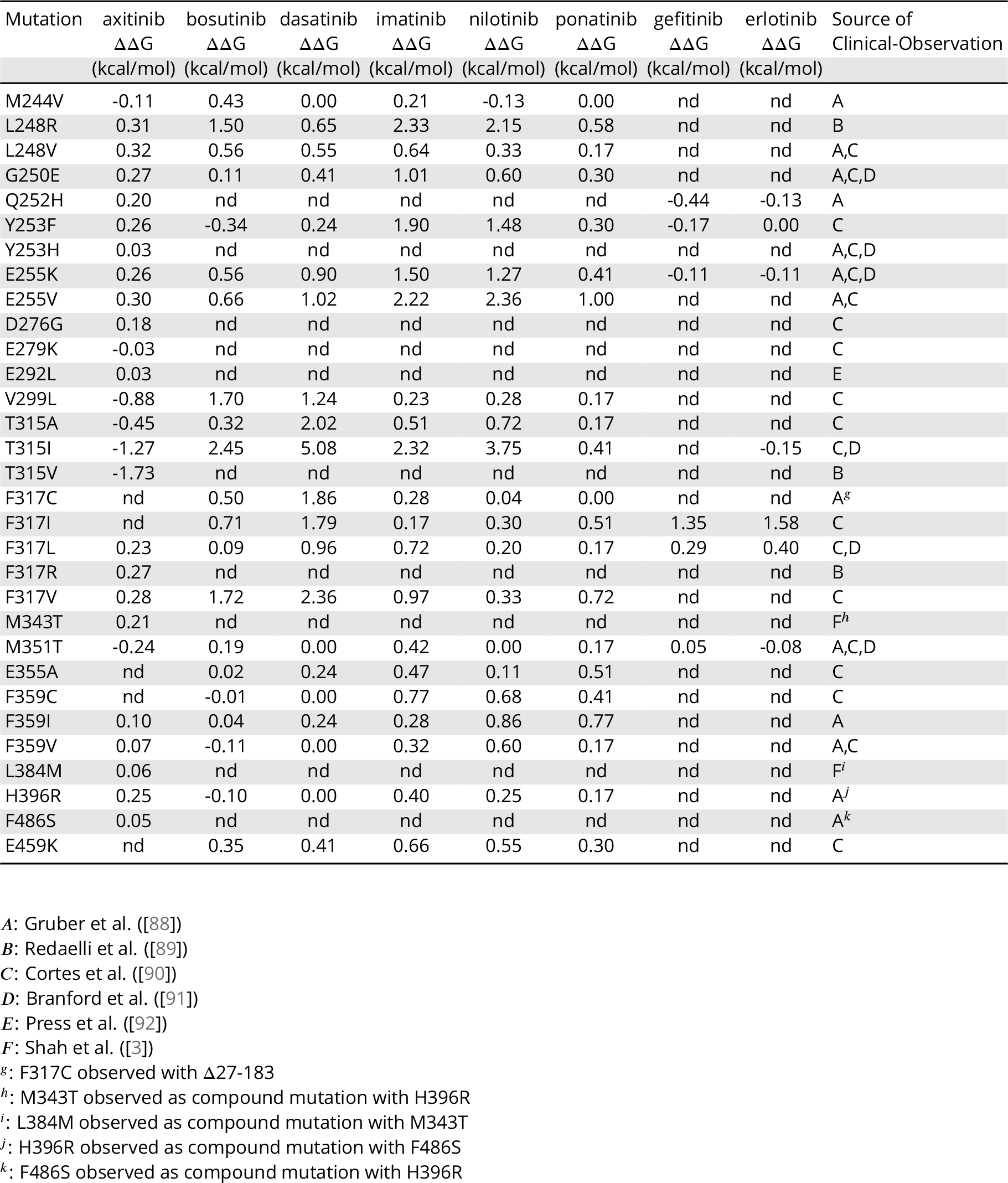
ΔΔG data derived from publicly available ΔpIC_50_ measurements and sources of mutation clinical-observation

**Table S2.** Axitinib: experimental IC_50_ values and alchemical free-energy ΔΔGs for each mutation.

| | Expt.<br>$IC_{50}$<br>(nM) | Expt.<br>$\Delta\Delta G$<br>(kcal/mol) | Prime<br>$\Delta\Delta G$<br>(kcal/mol) | FEP+<br>$\Delta\Delta G$<br>(kcal/mol) | FEP+<br>BAR err<br>(kcal/mol) | FEP+<br>$\Delta\Delta G$<br>(kcal/mol) | FEP+<br>BAR err<br>(kcal/mol) | FEP+<br>$\Delta\Delta G$<br>(kcal/mol) | FEP+<br>BAR err<br>(kcal/mol) | $\Delta\Delta G_{Av}$<br>(kcal/mol) | SE<br>(kcal/mol) |
| --- | --- | --- | --- | --- | --- | --- | --- | --- | --- | --- | --- |
| wild-type | 823 |  |  |  |  |  |  |  |  |  |  |
| M244V | 690 | -0.11 | -0.10 | -0.40 | 0.41 | -0.35 | 0.41 | -0.43 | 0.41 | -0.39 | 0.02 |
| L248R | 1393 | 0.31 | -0.06 | 2.13 | 0.43 | 2.42 | 0.45 | 2.46 | 0.43 | 2.34 | 0.10 |
| L248V | 1399 | 0.32 | 6.02 | -1.32 | 0.41 | -1.04 | 0.42 | -1.22 | 0.42 | -1.19 | 0.08 |
| G250E | 1295 | 0.27 | 0.31 | -0.35 | 0.41 | -0.71 | 0.41 | -0.74 | 0.41 | -0.60 | 0.13 |
| Q252H | 1155 | 0.20 | -0.18 | 0.07 | 0.43 | 0.30 | 0.42 | 0.29 | 0.43 | 0.22 | 0.08 |
| Y253F | 1275 | 0.26 | 1.11 | 0.77 | 0.43 | 0.23 | 0.43 | 1.15 | 0.45 | 0.72 | 0.27 |
| Y253H | 867 | 0.03 | 4.65 | 1.14 | 0.47 | 0.38 | 0.49 | -0.19 | 0.45 | 0.44 | 0.39 |
| E255K | 1282 | 0.26 | 0.12 | 1.30 | 0.44 | 0.63 | 0.43 | 1.10 | 0.44 | 1.01 | 0.20 |
| E255V | 1350 | 0.30 | -0.29 | 0.98 | 0.42 | 1.04 | 0.42 | 1.26 | 0.43 | 1.09 | 0.09 |
| D276G | 1105 | 0.18 | -0.01 | 0.03 | 0.42 | 0.64 | 0.42 | 0.44 | 0.43 | 0.37 | 0.18 |
| E279K | 778 | -0.03 | -0.15 | 0.06 | 0.42 | -0.22 | 0.43 | 1.27 | 0.43 | 0.37 | 0.46 |
| E292L | 863 | 0.03 | -0.00 | 0.53 | 0.43 | 0.35 | 0.42 | 0.31 | 0.42 | 0.40 | 0.07 |
| V299L | 188 | -0.88 | -5.00 | -1.08 | 0.42 | -1.39 | 0.42 | -1.37 | 0.42 | -1.28 | 0.10 |
| T315A | 389 | -0.45 | 0.99 | 0.09 | 0.43 | 0.24 | 0.47 | 0.31 | 0.42 | 0.21 | 0.06 |
| T315I | 98 | -1.27 | -2.30 | -1.26 | 0.42 | -1.50 | 0.45 | -1.39 | 0.43 | -1.38 | 0.07 |
| T315V | 45 | -1.73 | -1.07 | -1.10 | 0.41 | -1.32 | 0.42 | -1.15 | 0.48 | -1.19 | 0.07 |
| F317L | 1220 | 0.23 | 1.29 | -0.64 | 0.41 | -0.10 | 0.41 | -0.38 | 0.41 | -0.37 | 0.16 |
| F317R | 1286 | 0.27 | -2.46 | 2.64 | 0.46 | 2.27 | 0.51 | 1.38 | 0.47 | 2.10 | 0.37 |
| F317V | 1320 | 0.28 | 2.29 | 0.45 | 0.42 | 0.70 | 0.42 | 0.75 | 0.42 | 0.63 | 0.09 |
| M343T | 1175 | 0.21 | -0.04 | -0.26 | 0.54 | -0.50 | 0.53 | -0.58 | 0.50 | -0.45 | 0.10 |
| M351T | 553 | -0.24 | -0.07 | -0.25 | 0.41 | -0.03 | 0.41 | 0.37 | 0.41 | 0.03 | 0.18 |
| F359I | 975 | 0.10 | -0.04 | 1.89 | 0.41 | 1.60 | 0.42 | 1.78 | 0.41 | 1.76 | 0.08 |
| F359V | 933 | 0.07 | -0.07 | 2.68 | 0.42 | 1.55 | 0.42 | 1.64 | 0.41 | 1.96 | 0.36 |
| L384M | 916 | 0.06 | -0.01 | -0.07 | 0.41 | 0.27 | 0.41 | 0.23 | 0.41 | 0.14 | 0.11 |
| H396R | 1247 | 0.25 | -0.02 | 0.36 | 0.42 | 1.23 | 0.41 | 0.65 | 0.42 | 0.75 | 0.26 |
| F486S | 897 | 0.05 | -0.09 | 0.65 | 0.47 | 1.14 | 0.46 | 0.44 | 0.48 | 0.74 | 0.21 |

**BAR err:** Bennett Acceptance Ratio error. $\Delta\Delta G_{Av}$ : Average of three independent FEP+ runs.**SE:** Standard Error between three independent FEP+ runs.

**Table S3.** Bosutinib: experimental IC_50_ values and alchemical free-energy ΔΔGs for each mutation.

| | Expt.<br>IC<sub>50</sub><br>(nM) | Expt.<br>$\Delta\Delta G$<br>(kcal/mol) | Prime<br>$\Delta\Delta G$<br>(kcal/mol) | FEP+<sub>Run1</sub><br>$\Delta\Delta G$<br>(kcal/mol) | FEP+<sub>Run1</sub><br>BAR err<br>(kcal/mol) | FEP+<sub>Run2</sub><br>$\Delta\Delta G$<br>(kcal/mol) | FEP+<sub>Run2</sub><br>BAR err<br>(kcal/mol) | FEP+<sub>Run3</sub><br>$\Delta\Delta G$<br>(kcal/mol) | FEP+<sub>Run3</sub><br>BAR err<br>(kcal/mol) | $\Delta\Delta G_{Av}$<br>(kcal/mol) | SE<br>(kcal/mol) |
| --- | --- | --- | --- | --- | --- | --- | --- | --- | --- | --- | --- |
| wild-type | 71 |  |  |  |  |  |  |  |  |  |  |
| M244V | 147 | 0.43 | 0.02 | -0.28 | 0.41 | -0.11 | 0.41 | -0.08 | 0.41 | -0.16 | 0.06 |
| L248R | 874 | 1.50 | 3.67 | 1.00 | 0.43 | 1.63 | 0.43 | 1.33 | 0.43 | 1.32 | 0.18 |
| L248V | 182 | 0.56 | 5.77 | 0.37 | 0.41 | 0.72 | 0.42 | 0.38 | 0.42 | 0.49 | 0.12 |
| G250E | 85 | 0.11 | -0.30 | 0.28 | 0.43 | 0.63 | 0.43 | -1.07 | 0.43 | -0.05 | 0.52 |
| Y253F | 40 | -0.34 | -0.03 | 0.21 | 0.45 | 0.02 | 0.43 | 0.95 | 0.43 | 0.39 | 0.28 |
| E255K | 181 | 0.56 | 0.49 | -1.01 | 0.43 | -1.30 | 0.43 | -1.01 | 0.43 | -1.11 | 0.10 |
| E255V | 214 | 0.66 | 0.11 | -0.47 | 0.42 | -0.51 | 0.43 | -0.91 | 0.43 | -0.63 | 0.14 |
| V299L | 1228 | 1.70 | -0.85 | 0.97 | 0.43 | 0.90 | 0.42 | 0.85 | 0.42 | 0.91 | 0.03 |
| T315A | 122 | 0.32 | 1.00 | -1.61 | 0.41 | -1.61 | 0.41 | -1.97 | 0.41 | -1.73 | 0.12 |
| T315I | 4338 | 2.45 | 3.75 | -2.32 | 0.43 | -2.21 | 0.42 | -1.26 | 0.42 | -1.93 | 0.34 |
| F317C | 165 | 0.50 | 4.83 | 1.04 | 0.41 | 1.27 | 0.41 | 1.22 | 0.42 | 1.18 | 0.07 |
| F317I | 232 | 0.71 | 1.61 | 0.16 | 0.41 | 0.07 | 0.42 | 0.02 | 0.41 | 0.08 | 0.04 |
| F317L | 82 | 0.09 | -0.71 | 0.05 | 0.41 | 0.47 | 0.41 | 0.24 | 0.41 | 0.25 | 0.12 |
| F317V | 1280 | 1.72 | 4.12 | 1.98 | 0.42 | 1.50 | 0.42 | 2.25 | 0.42 | 1.91 | 0.22 |
| M351T | 97 | 0.19 | 0.02 | 0.36 | 0.42 | 0.82 | 0.41 | 0.71 | 0.41 | 0.63 | 0.14 |
| E355A | 74 | 0.02 | 0.13 | -0.20 | 0.44 | 0.13 | 0.43 | 0.27 | 0.43 | 0.07 | 0.14 |
| F359C | 70 | -0.01 | -0.09 | 3.02 | 0.42 | 2.51 | 0.42 | 1.97 | 0.43 | 2.50 | 0.30 |
| F359I | 76 | 0.04 | -0.06 | 0.66 | 0.41 | 1.74 | 0.41 | 1.43 | 0.42 | 1.28 | 0.32 |
| F359V | 59 | -0.11 | -0.06 | 0.98 | 0.43 | 1.69 | 0.41 | 1.91 | 0.42 | 1.53 | 0.28 |
| H396R | 60 | -0.10 | -1.07 | 0.62 | 0.42 | -0.07 | 0.42 | -0.93 | 0.43 | -0.13 | 0.45 |
| E459K | 127 | 0.35 | 0.26 | -0.69 | 0.42 | 0.23 | 0.42 | -0.54 | 0.42 | -0.33 | 0.28 |

**BAR err:** Bennett Acceptance Ratio error. $\Delta\Delta G_{Av}$ : Average of three independent FEP+ runs.**SE:** Standard Error between three independent FEP+ runs.

**Table S4.** Dasatinib: experimental IC_50_ values and alchemical free-energy ΔΔGs for each mutation.

| | Expt.<br>IC<sub>50</sub><br>(nM) | Expt.<br>$\Delta\Delta G$<br>(kcal/mol) | Prime<br>$\Delta\Delta G$<br>(kcal/mol) | FEP+<sub>Run1</sub><br>$\Delta\Delta G$<br>(kcal/mol) | FEP+<sub>Run1</sub><br>BAR err<br>(kcal/mol) | FEP+<sub>Run2</sub><br>$\Delta\Delta G$<br>(kcal/mol) | FEP+<sub>Run2</sub><br>BAR err<br>(kcal/mol) | FEP+<sub>Run3</sub><br>$\Delta\Delta G$<br>(kcal/mol) | FEP+<sub>Run3</sub><br>BAR err<br>(kcal/mol) | $\Delta\Delta G_{Av}$<br>(kcal/mol) | SE<br>(kcal/mol) |
| --- | --- | --- | --- | --- | --- | --- | --- | --- | --- | --- | --- |
| wild-type | 2 |  |  |  |  |  |  |  |  |  |  |
| M244V | 2 | 0.00 | -0.10 | 0.05 | 0.41 | -0.37 | 0.41 | -0.43 | 0.41 | -0.25 | 0.15 |
| L248R | 6 | 0.65 | -2.13 | 1.40 | 0.42 | 1.50 | 0.43 | 1.51 | 0.42 | 1.47 | 0.04 |
| L248V | 5 | 0.55 | 2.60 | 0.58 | 0.42 | 0.70 | 0.41 | 0.79 | 0.41 | 0.69 | 0.06 |
| G250E | 4 | 0.41 | -0.00 | -0.54 | 0.43 | -0.31 | 0.43 | 0.01 | 0.44 | -0.28 | 0.16 |
| Y253F | 3 | 0.24 | 0.00 | -0.21 | 0.43 | -0.24 | 0.43 | -0.03 | 0.44 | -0.16 | 0.07 |
| E255K | 9 | 0.90 | -0.08 | -0.30 | 0.43 | -0.17 | 0.44 | -1.05 | 0.43 | -0.51 | 0.27 |
| E255V | 11 | 1.02 | -0.08 | 0.06 | 0.42 | -0.80 | 0.42 | -0.12 | 0.42 | -0.29 | 0.26 |
| V299L | 16 | 1.24 | 0.01 | 0.83 | 0.41 | 0.36 | 0.42 | 0.77 | 0.42 | 0.65 | 0.15 |
| T315A | 59 | 2.02 | 5.09 | -1.74 | 0.41 | -1.65 | 0.41 | -1.23 | 0.41 | -1.54 | 0.16 |
| T315I | 10000 | 5.08 | -2.69 | 5.63 | 0.43 | 4.69 | 0.44 | 5.50 | 0.43 | 5.27 | 0.29 |
| F317C | 45 | 1.86 | 4.72 | 2.63 | 0.42 | 2.32 | 0.42 | 2.62 | 0.41 | 2.52 | 0.10 |
| F317I | 40 | 1.79 | 2.38 | 1.94 | 0.41 | 2.04 | 0.41 | 1.94 | 0.41 | 1.97 | 0.03 |
| F317L | 10 | 0.96 | 1.22 | 1.26 | 0.41 | 1.42 | 0.41 | 1.08 | 0.41 | 1.25 | 0.10 |
| F317V | 104 | 2.36 | 4.08 | 3.12 | 0.42 | 2.84 | 0.42 | 2.68 | 0.42 | 2.88 | 0.13 |
| M351T | 2 | 0.00 | 0.04 | 0.04 | 0.41 | 0.14 | 0.41 | 0.00 | 0.42 | 0.06 | 0.04 |
| E355A | 3 | 0.24 | 0.00 | -0.24 | 0.43 | -0.87 | 0.45 | -1.25 | 0.44 | -0.79 | 0.29 |
| F359C | 2 | 0.00 | -0.03 | 1.24 | 0.42 | 0.68 | 0.41 | 1.38 | 0.42 | 1.10 | 0.21 |
| F359I | 3 | 0.24 | -0.02 | -0.50 | 0.42 | -0.33 | 0.42 | -1.14 | 0.42 | -0.66 | 0.25 |
| F359V | 2 | 0.00 | -0.03 | -0.87 | 0.41 | 0.57 | 0.42 | -0.62 | 0.41 | -0.31 | 0.44 |
| H396R | 2 | 0.00 | 2.53 | -0.76 | 0.43 | -0.09 | 0.43 | -0.06 | 0.43 | -0.30 | 0.23 |
| E459K | 4 | 0.41 | 0.00 | -0.68 | 0.42 | -0.17 | 0.42 | -0.07 | 0.41 | -0.31 | 0.19 |

T315I was beyond the concentration limit of the assay (10,000 nM).
**BAR err:** Bennett Acceptance Ratio error.
$\Delta\Delta G_{Av}$ : Average of three independent FEP+ runs.
**SE:** Standard Error between three independent FEP+ runs.

**Table S5.** Imatinib: experimental IC_50_ values and alchemical free-energy ΔΔGs for each mutation.

| | Expt.<br>IC<sub>50</sub><br>(nM) | Expt.<br>$\Delta\Delta G$<br>(kcal/mol) | Prime<br>$\Delta\Delta G$<br>(kcal/mol) | FEP+<sub>Run1</sub><br>$\Delta\Delta G$<br>(kcal/mol) | FEP+<sub>Run1</sub><br>BAR err<br>(kcal/mol) | FEP+<sub>Run2</sub><br>$\Delta\Delta G$<br>(kcal/mol) | FEP+<sub>Run2</sub><br>BAR err<br>(kcal/mol) | FEP+<sub>Run3</sub><br>$\Delta\Delta G$<br>(kcal/mol) | FEP+<sub>Run3</sub><br>BAR err<br>(kcal/mol) | $\Delta\Delta G_{Av}$<br>(kcal/mol) | SE<br>(kcal/mol) |
| --- | --- | --- | --- | --- | --- | --- | --- | --- | --- | --- | --- |
| wild-type | 201 |  |  |  |  |  |  |  |  |  |  |
| M244V | 287 | 0.21 | -0.08 | 0.15 | 0.41 | 0.43 | 0.41 | 0.17 | 0.41 | 0.25 | 0.09 |
| L248R | 10000 | 2.33 | 1.92 | 1.92 | 0.43 | 2.52 | 0.44 | 2.34 | 0.43 | 2.26 | 0.18 |
| L248V | 586 | 0.64 | 1.89 | -1.04 | 0.41 | -1.02 | 0.42 | -1.20 | 0.41 | -1.09 | 0.06 |
| G250E | 1087 | 1.01 | 0.92 | 0.16 | 0.41 | 0.02 | 0.41 | 0.12 | 0.41 | 0.10 | 0.04 |
| Y253F | 4908 | 1.90 | -0.02 | 0.87 | 0.43 | 0.65 | 0.42 | 1.34 | 0.44 | 0.95 | 0.20 |
| E255K | 2487 | 1.50 | 0.25 | -0.12 | 0.44 | 1.95 | 0.44 | -0.55 | 0.44 | 0.43 | 0.77 |
| E255V | 8322 | 2.22 | 0.24 | -0.72 | 0.42 | -0.02 | 0.42 | -0.53 | 0.43 | -0.42 | 0.21 |
| V299L | 295 | 0.23 | -1.29 | 0.66 | 0.41 | 0.26 | 0.42 | -0.37 | 0.42 | 0.18 | 0.30 |
| T315A | 476 | 0.51 | 5.10 | -1.39 | 0.41 | -1.86 | 0.41 | -2.09 | 0.44 | -1.78 | 0.21 |
| T315I | 9773 | 2.32 | 0.88 | 4.23 | 0.43 | 4.23 | 0.42 | 3.14 | 0.44 | 3.87 | 0.36 |
| F317C | 324 | 0.28 | 2.10 | 0.27 | 0.42 | -0.18 | 0.41 | 0.45 | 0.42 | 0.18 | 0.19 |
| F317I | 266 | 0.17 | 0.94 | 0.59 | 0.41 | 0.66 | 0.41 | 0.48 | 0.41 | 0.58 | 0.05 |
| F317L | 675 | 0.72 | 0.74 | 0.58 | 0.41 | 0.53 | 0.41 | 0.38 | 0.41 | 0.50 | 0.06 |
| F317V | 1023 | 0.97 | 1.57 | 0.71 | 0.42 | 0.79 | 0.42 | 0.80 | 0.41 | 0.77 | 0.03 |
| M351T | 404 | 0.42 | -0.02 | 1.72 | 0.41 | 1.03 | 0.42 | 1.20 | 0.42 | 1.32 | 0.21 |
| E355A | 441 | 0.47 | 0.29 | 0.13 | 0.43 | 0.08 | 0.44 | 0.14 | 0.43 | 0.12 | 0.02 |
| F359C | 728 | 0.77 | 2.43 | 0.88 | 0.42 | 0.47 | 0.41 | 0.33 | 0.42 | 0.56 | 0.17 |
| F359I | 324 | 0.28 | 1.95 | -0.13 | 0.41 | -0.87 | 0.41 | 0.08 | 0.41 | -0.31 | 0.29 |
| F359V | 346 | 0.32 | 2.53 | -0.66 | 0.41 | 0.02 | 0.41 | -0.27 | 0.42 | -0.30 | 0.20 |
| H396R | 395 | 0.40 | 2.76 | -0.39 | 0.41 | -0.38 | 0.42 | -0.39 | 0.42 | -0.39 | 0.00 |
| E459K | 612 | 0.66 | 0.24 | -0.09 | 0.43 | -0.09 | 0.42 | -0.08 | 0.42 | -0.09 | 0.00 |

T315I was beyond the concentration limit of the assay (10,000 nM).
**BAR err:** Bennett Acceptance Ratio error.
$\Delta\Delta G_{Av}$ : Average of three independent FEP+ runs.
**SE:** Standard Error between three independent FEP+ runs.

**Table S6.** Nilotinib: experimental IC_50_ values and alchemical free-energy ΔΔGs for each mutation.

| | Expt.<br>$IC_{50}$<br>(nM) | Expt.<br>$\Delta\Delta G$<br>(kcal/mol) | Prime<br>$\Delta\Delta G$<br>(kcal/mol) | FEP+<sub>Run1</sub><br>$\Delta\Delta G$<br>(kcal/mol) | FEP+<sub>Run1</sub><br>BAR err<br>(kcal/mol) | FEP+<sub>Run2</sub><br>$\Delta\Delta G$<br>(kcal/mol) | FEP+<sub>Run2</sub><br>BAR err<br>(kcal/mol) | FEP+<sub>Run3</sub><br>$\Delta\Delta G$<br>(kcal/mol) | FEP+<sub>Run3</sub><br>BAR err<br>(kcal/mol) | $\Delta\Delta G_{Av}$<br>(kcal/mol) | SE<br>(kcal/mol) |
| --- | --- | --- | --- | --- | --- | --- | --- | --- | --- | --- | --- |
| wild-type | 15 |  |  |  |  |  |  |  |  |  |  |
| M244V | 12 | -0.13 | -0.11 | 0.15 | 0.41 | -0.21 | 0.41 | 0.21 | 0.41 | 0.05 | 0.13 |
| L248R | 549 | 2.15 | 0.48 | 2.05 | 0.43 | 2.12 | 0.47 | 1.93 | 0.43 | 2.03 | 0.06 |
| L248V | 26 | 0.33 | 3.53 | -0.50 | 0.42 | -0.39 | 0.41 | -0.92 | 0.41 | -0.60 | 0.16 |
| G250E | 41 | 0.60 | 0.05 | 0.06 | 0.41 | -0.27 | 0.41 | -0.38 | 0.41 | -0.20 | 0.13 |
| Y253F | 179 | 1.48 | -0.27 | 1.09 | 0.43 | 0.42 | 0.42 | 1.16 | 0.42 | 0.89 | 0.24 |
| E255K | 127 | 1.27 | 0.41 | -2.24 | 0.48 | -1.52 | 0.46 | 0.33 | 0.46 | -1.14 | 0.77 |
| E255V | 784 | 2.36 | -0.03 | 0.31 | 0.42 | -0.25 | 0.43 | -0.55 | 0.43 | -0.16 | 0.25 |
| V299L | 24 | 0.28 | 2.94 | -0.18 | 0.41 | 0.21 | 0.41 | 0.15 | 0.41 | 0.06 | 0.12 |
| T315A | 50 | 0.72 | 3.38 | -1.33 | 0.41 | -1.31 | 0.41 | -1.39 | 0.41 | -1.34 | 0.02 |
| T315I | 8091 | 3.75 | 4.16 | 4.29 | 0.43 | 5.00 | 0.42 | 4.34 | 0.43 | 4.54 | 0.23 |
| F317C | 16 | 0.04 | 0.90 | 1.34 | 0.41 | 0.88 | 0.41 | 0.60 | 0.41 | 0.94 | 0.22 |
| F317I | 25 | 0.30 | -0.18 | 1.24 | 0.41 | 1.17 | 0.41 | 0.82 | 0.41 | 1.08 | 0.13 |
| F317L | 21 | 0.20 | 1.74 | 1.03 | 0.41 | 1.07 | 0.41 | 1.09 | 0.41 | 1.06 | 0.02 |
| F317V | 26 | 0.33 | 0.77 | 1.16 | 0.41 | 0.68 | 0.42 | 1.07 | 0.42 | 0.97 | 0.15 |
| M351T | 15 | 0.00 | 0.09 | -0.06 | 0.41 | -0.09 | 0.42 | -0.46 | 0.42 | -0.20 | 0.13 |
| E355A | 18 | 0.11 | -0.06 | -0.46 | 0.43 | -1.01 | 0.43 | -0.32 | 0.43 | -0.60 | 0.21 |
| F359C | 47 | 0.68 | 3.68 | 1.32 | 0.41 | 1.44 | 0.41 | 1.52 | 0.41 | 1.43 | 0.06 |
| F359I | 64 | 0.86 | 3.70 | 1.05 | 0.41 | 1.13 | 0.41 | 0.74 | 0.41 | 0.97 | 0.12 |
| F359V | 41 | 0.60 | 3.67 | 1.00 | 0.41 | 1.08 | 0.41 | 1.38 | 0.42 | 1.15 | 0.12 |
| H396R | 23 | 0.25 | 2.58 | -0.07 | 0.42 | 0.21 | 0.42 | 0.03 | 0.42 | 0.06 | 0.08 |
| E459K | 38 | 0.55 | -0.00 | -0.17 | 0.42 | -0.46 | 0.42 | -0.10 | 0.42 | -0.24 | 0.11 |

**BAR err:** Bennett Acceptance Ratio error. $\Delta\Delta G_{Av}$ : Average of three independent FEP+ runs.**SE:** Standard Error between three independent FEP+ runs.

**Table S7.** Ponatinib: experimental IC_50_ values and alchemical free-energy ΔΔGs for each mutation.

| | Expt.<br>IC<sub>50</sub><br>(nM) | Expt.<br>$\Delta\Delta$G<br>(kcal/mol) | Prime<br>$\Delta\Delta$G<br>(kcal/mol) | FEP+<sub>Run1</sub><br>$\Delta\Delta$G<br>(kcal/mol) | FEP+<sub>Run1</sub><br>BAR err<br>(kcal/mol) | FEP+<sub>Run2</sub><br>$\Delta\Delta$G<br>(kcal/mol) | FEP+<sub>Run2</sub><br>BAR err<br>(kcal/mol) | FEP+<sub>Run3</sub><br>$\Delta\Delta$G<br>(kcal/mol) | FEP+<sub>Run3</sub><br>BAR err<br>(kcal/mol) | $\Delta\Delta$G<sub>Av</sub><br>(kcal/mol) | SE<br>(kcal/mol) |
| --- | --- | --- | --- | --- | --- | --- | --- | --- | --- | --- | --- |
| wild-type | 3 |  |  |  |  |  |  |  |  |  |  |
| M244V | 3 | 0.00 | -0.13 | 0.07 | 0.41 | -0.28 | 0.41 | 0.12 | 0.41 | -0.03 | 0.13 |
| L248R | 8 | 0.58 | 2.48 | 1.40 | 0.43 | 0.96 | 0.43 | 1.10 | 0.44 | 1.15 | 0.13 |
| L248V | 4 | 0.17 | 2.48 | -1.82 | 0.42 | -1.23 | 0.42 | -1.96 | 0.42 | -1.67 | 0.22 |
| G250E | 0.021 | 0.30 | 0.17 | -0.32 | 0.43 | -0.25 | 0.43 | -0.71 | 0.46 | -0.43 | 0.14 |
| Y253F | 5 | 0.30 | 0.05 | 0.85 | 0.43 | 1.32 | 0.44 | 0.77 | 0.43 | 0.98 | 0.17 |
| E255K | 6 | 0.41 | 1.05 | -0.27 | 0.48 | -0.66 | 0.48 | 0.03 | 0.47 | -0.30 | 0.20 |
| E255V | 16 | 1.00 | -0.04 | 1.19 | 0.43 | 0.94 | 0.43 | -0.41 | 0.43 | 0.57 | 0.50 |
| V299L | 4 | 0.17 | -0.29 | -0.56 | 0.41 | -0.55 | 0.41 | -1.42 | 0.41 | -0.84 | 0.29 |
| T315A | 4 | 0.17 | -0.51 | -2.90 | 0.41 | -3.15 | 0.41 | -2.92 | 0.41 | -2.99 | 0.08 |
| T315I | 6 | 0.41 | -5.42 | 0.51 | 0.42 | 0.90 | 0.42 | 0.91 | 0.42 | 0.77 | 0.13 |
| F317C | 3 | 0.00 | 1.45 | 0.44 | 0.41 | 0.98 | 0.42 | 0.80 | 0.41 | 0.74 | 0.16 |
| F317I | 7 | 0.51 | 0.62 | -0.76 | 0.41 | -1.03 | 0.41 | -1.02 | 0.41 | -0.94 | 0.09 |
| F317L | 4 | 0.17 | 0.57 | -1.08 | 0.41 | -0.83 | 0.41 | -0.85 | 0.41 | -0.92 | 0.08 |
| F317V | 10 | 0.72 | 1.14 | 0.05 | 0.41 | -0.21 | 0.41 | 0.24 | 0.42 | 0.03 | 0.13 |
| M351T | 4 | 0.17 | -0.12 | 0.89 | 0.41 | 1.66 | 0.41 | 0.65 | 0.41 | 1.07 | 0.30 |
| E355A | 7 | 0.51 | 0.01 | 0.12 | 0.44 | -0.52 | 0.44 | -0.55 | 0.43 | -0.32 | 0.22 |
| F359C | 6 | 0.41 | 2.12 | 0.25 | 0.42 | -0.35 | 0.43 | 0.73 | 0.42 | 0.21 | 0.31 |
| F359I | 11 | 0.77 | 0.34 | -0.66 | 0.41 | -0.38 | 0.41 | 0.06 | 0.41 | -0.33 | 0.21 |
| F359V | 4 | 0.17 | 0.74 | 0.11 | 0.41 | -0.28 | 0.41 | 0.08 | 0.42 | -0.03 | 0.13 |
| H396R | 4 | 0.17 | -0.04 | 0.19 | 0.49 | 0.10 | 0.45 | -1.41 | 0.48 | -0.37 | 0.52 |
| E459K | 5 | 0.30 | -0.00 | -0.51 | 0.42 | -0.78 | 0.42 | -0.63 | 0.42 | -0.64 | 0.08 |

**Table S8.**
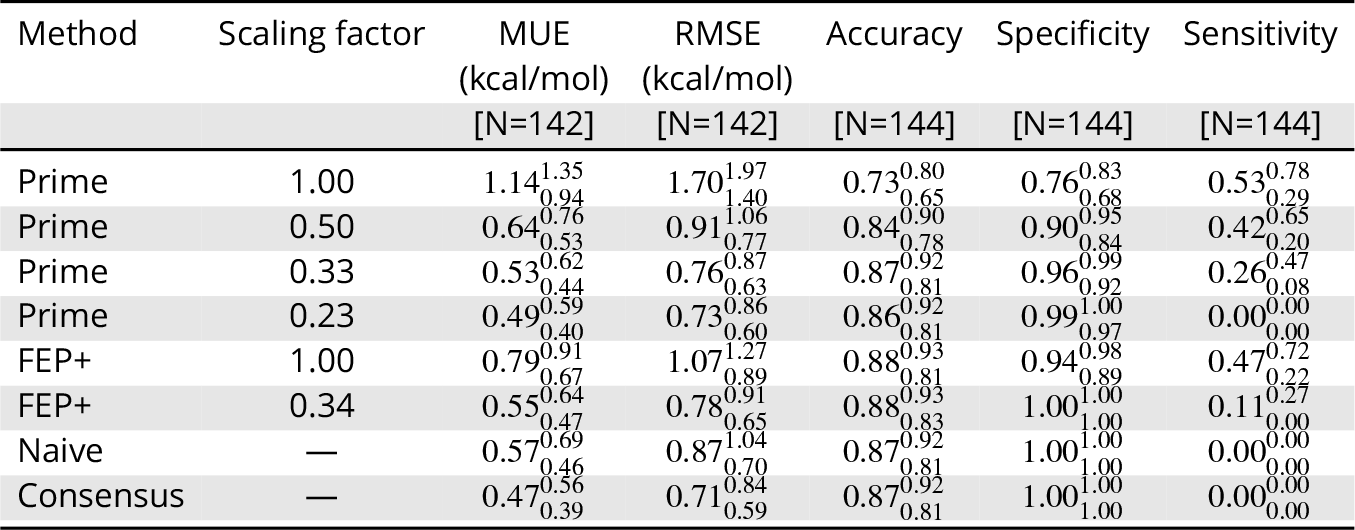
Summary of statistics of scaled predictions, a naïve model, and a consensus model.

**Table S9.**
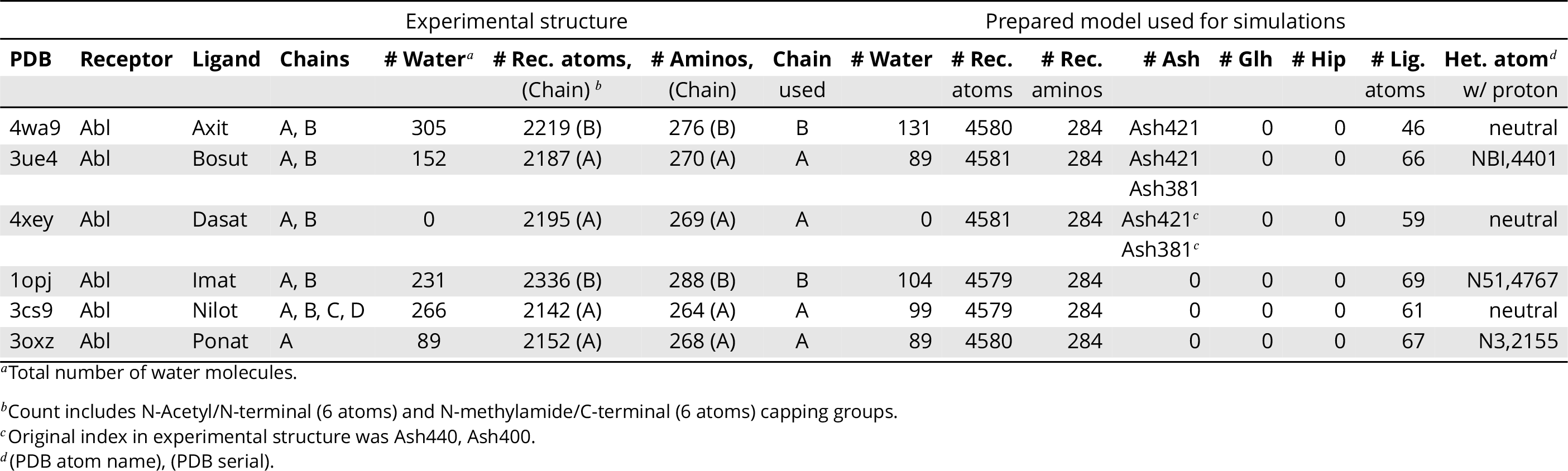
Summary of the preparation of the 6 Abl:TKI co-crystal structure complexes.

